# Structural basis for ACT1 oligomerization induced by IL-17 receptor hetero-tetramer

**DOI:** 10.64898/2025.12.02.691943

**Authors:** Hui Zhang, Xiao-chen Bai, Xuewu Zhang

**Affiliations:** Department of Pharmacology, University of Texas Southwestern Medical Center, Dallas, Texas, USA; Department of Biophysics, University of Texas Southwestern Medical Center, Dallas, Texas, USA; Department of Cell Biology, University of Texas Southwestern Medical Center, Dallas, Texas, USA

**Author notes:** Corresponding authors (X-C. B.); (X.Z.).

## Abstract

The IL-17 receptors (IL-17Rs) play critical roles in immunity and inflammatory diseases. IL-17-induced heteromeric complexes between IL-17RA and another IL-17R trigger signaling by binding the downstream transducer ACT1 through interactions between their intracellular SEF/IL-17R (SEFIR) domains. The molecular mechanism of this process remains unclear. Here we present the cryo-EM structure of the complex of IL-17RA, IL-17RB and ACT1, showing that the IL-17RA and IL-17RB SEFIR domains form an asymmetric hetero-tetramer. The two IL-17RA SEFIR domains serve as the base to recruit ACT1, while IL-17RB stabilizes the IL-17RA dimer but makes no interaction with ACT1. IL-17RB, IL-17RA and multiple ACT1 together form a double-stranded helical assembly. The C-terminal SEFIR extension (SEFEX) of IL-17RA acts as a molecular tendril to help anchor the ACT1 protomers. The structural model is supported by our mutational analyses. These findings reveal the basis for the formation for the signalosome of the IL-17 receptors and ACT1 critical for immune signaling.

## Introduction

The IL-17 family cytokines (IL-17A, B, C, D, E and F) are proinflammatory cytokines secreted by Th17 cells, innate immune cells as well as non-immune cell types that play important roles in immune defense against infection, tissue barrier protection and repair by activating proinflammatory proteins such as NF-kB^1–4^. Loss of IL-17 signaling due to mutations of IL-17, their receptors or downstream signaling proteins cause defects in immune responses against bacterial and fungal infection^3^. Conversely, chronical IL-17 signaling can promote cancer, inflammatory and autoimmune diseases such as psoriasis, arthritis and asthma^3,4^. Antibodies that target IL-17 and their receptors are widely used in clinic for treating these diseases^4,5^. More recent studies have shown that IL-17 signaling in the brain can regulate mood and anxiety, providing an explanation for mood alterations in patients suffering from autoimmune or infectious diseases^6,7^.

The IL-17 cytokines function as homodimers or heterodimers that activate the IL-17 family receptors (IL-17Rs), which contain five members (RA, RB, RC, RD and RE)^8^. The IL-17Rs are single-pass transmembrane proteins that use their extracellular region to engage IL-17, while the intracellular region transduces the signal to downstream pathways (Fig. 1a). Several studies have shown that heteromeric complexes formed by IL-17RA and another IL-17 receptor family member are required for transducing IL-17 signal^9,10^, although one study has suggested that IL-17RC can function as a homodimer for IL-17RA-independent signaling^11^. Recent structural studies have shown that different IL-17 dimers induce active IL-17 receptor assemblies through different binding modes^12,13^. For example, the homodimer of IL-17E (also known as IL-25) binds two IL-17RB molecules, inducing a conformational change in IL-17RB that allows it to recruit IL-17RA, leading to a 2:2:2 IL-17E/IL-17RA/IL-17RB assembly ^13^. On the other hand, the IL-17A dimer induces the formation of the complex containing one IL-17RA and one IL-17RC^12,13^. Two copies of this complex further assemble into the functional 4:2:2 complex through an interaction between the two IL-17RA molecules. Regardless the binding modes, it appears that the signaling requires the assembly of two IL-17RA molecules act as the common receptor and two protomers of another IL-17 receptor family member act as the co-receptor^12,13^.

**Fig 1.**
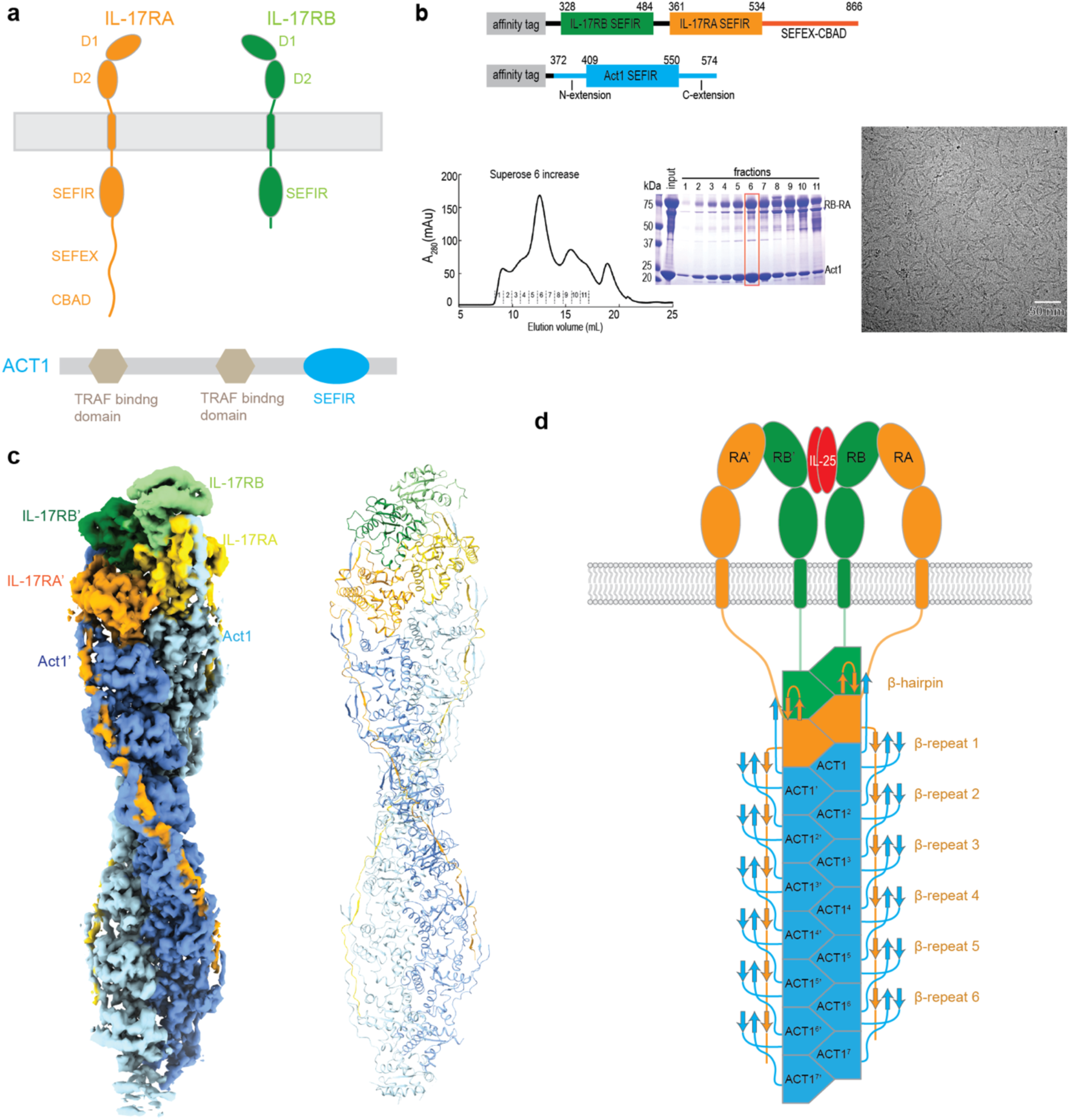
Cryo-EM structure of the IL-17RA/IL-17RB/ACT1 complex. **a**, Schematic domain organization of IL-17RA, IL-17RB, and ACT1. **b**, Expression and purification of the complex between the IL-17RB-RA fusion and ACT1. The upper panel shows the constructs for protein expression. The lower left and middle panels show the gel filtration chromatography profile and SDS-PAGE gel analyses of the complex. The peak fraction highlighted by the red box was used as the cryo-EM sample. The results shown are representatives of at least three repeats. A sample image of the IL-17RA/RB/ACT1 filaments is shown in the lower right panel. **c**, Cryo-EM map (left) and model (right) of the IL-17RA/RB/ACT1 complex. **d**, Cartoon representation of the structure of the IL-17RA/RB/ACT1 complex.

The IL-17/IL-17R complexes transduce signal to the inside of the cell through the intracellular SEFIR domain, a small domain related to the Toll/IL-IR (TIR) domains found in Toll/IL-1 family receptors (Fig. 1a)^8,14,15^. The key downstream signal transducer of IL-17R is ACT1 (NF-kB activator 1; also known as CIKS and TRFA3IP2), which also contains a SEFIR domain that interacts with the receptors by homotypic interactions (Fig. 1a)^16,17^. Besides the SEFIR domain, ACT1 contains several other functionally important elements in its N-terminal regions, which transduces signal by interacting with downstream proteins such as TRAF6 or functioning as an E3 ubiquitin ligase^8,18^. IL-17/IL-17R complexes promote ACT1 activation by inducing its oligomerization, leading to the formation of large signalosomes at the plasma membrane^12,19^. In addition to the SEFIR domain, the long C-terminal extension in IL-17RA contains a SEFIR-extension (SEFEX) and C/EBP-β activation domain (CBAD) (Fig. 1a)^8^. The SEFEX region plays an important role in binding and activating ACT1^20^. Other IL-17 receptors such as IL-17RB and IL-17RC do not contain a SEFEX, which may be the reason that IL-17RA is an obligate common receptor for IL-17 signaling.

Despite extensive investigations, it remains unclear how IL-17-activated IL-17Rs bind ACT1, why heteromeric IL-17R complexes are required for activating ACT1, and how ACT1 undergoes oligomerization for signaling. To address these questions, we solved the cryo-EM structure of the complex of the intracellular domains of IL-17RA, IL-17RB and ACT1, revealing a double-stranded helical assembly that contains high-order oligomers of ACT1. The structure and the associated functional assays elucidate the intricate interactions in the complex mediated by distinct structural features in the three proteins that underlie the formation of the IL-17/IL-17R/ACT1 signalosome.

## Results

### Structure determination of the complex of IL-17RA, IL-17RB and ACT1

To solve the structure of the intracellular domain complexes of *human* IL-17Rs and ACT1, we tried several strategies for assembling the complexes, including mixing separately purified various constructs of the SEFIR domains of IL-17Rs and ACT1 as well as co-expressing them in bacteria, insect or HEK293 cells. These efforts all failed, likely due to weak interactions between the individual proteins. Based on recent structural studies, the active IL-17R complexes induced by IL-17 are minimally hetero-tetrameric assemblies of two protomers of IL-17RA and two protomers of a co-receptor^12,13^. To facilitate the formation of the IL-17R SEFIR domain hetero-tetramers, we made a fusion construct between IL-17RB (residues 328-484) and IL-17RA (residues 361-866) with a 16-residue flexible linker between them (See method for details) (Fig. 1b). The IL-17RA part spans both the SEFIR domain and the SEFEX required for optimal activation of ACT1 in cells^20^. Co-expression of the IL-17RB-RA fusion with the construct of ACT1 spanning residues 372-574 (containing both the SEFIR domain and the conserved N- and C-terminal extensions) yielded a large stable complex with super-stoichiometric ACT1 as indicated by the gel filtration chromatography and SDS-gel results (Fig. 1b).

Cryo-EM images of the IL-17RA/RB/ACT1 complex showed abundant filamentous assemblies of 300-400 Å in length with an obvious helical twist (Fig. 1b). We solved the structure of this complex to 3.0 Å resolution by single-particle reconstruction, and built a model containing 2, 2 and 15 protomers of IL-17RA, IL17RB and ACT1, respectively (Fig. 1c and Supplementary Fig.1-3). Weak density at the end beyond the atomic model indicates that additional ACT1 subunits were present in a subset of particles. On the other hand, the presence of shorter filaments in micrograph shows that some filaments contain fewer ACT1 subunits. These observations together suggest that the IL-17R hetero-tetramers could promote the oligomerization of ACT1, although the length of the oligomer varies to some extent. Nevertheless, our atomic model captures the essential structure features that underlie the assembly of the IL17RA/RB/ACT1 signalosome.

### Overall structure of the IL-17RA/RB/ACT1 complex

The IL-17RA/RB/ACT1 complex forms a double-stranded filamentous assembly (Fig. 1c), reminiscent of those formed by related TIR domains of MyD88 (myeloid differentiation factor 88), Mal (MyD88 adaptor-like), TRIF (TIR-domain-containing adapter-inducing interferon- β) and TRIF (TRIF-related adaptor molecule) (Supplementary fig. 4a)^21–23^. However, unlike the straight overall shape of the protofilaments in those TIR domain filaments, the IL-17RA/RB/ACT1 complex displays a left-handed helical twist, with consecutive subunits in each protofilament related by a 33.6 Å shift and -34.2° twist along the axis of the double-helix (Supplementary Fig. 1). The two strands are related to each other by an axial offset of 16.8 Å, leading to staggered arrangement of the two subunits in each layer of the double helix. This arrangement, which is also present in the TIR domain structures, allows each subunit in the middle of the filament to interact with four other subunits, two from the same protofilament and two from the other protofilament. Notably, IL-17RA, RB and ACT1 have several structural elements that are distinct from one another and from the TIR domains, which form unique interactions in the complex. The most notable are the repetitive β-sheet modules formed by the long SEFEX region of IL-17RA and both the N- and C-terminal extensions of the ACT1 SEFIR domain (Fig. 1c and 1d). These features play important roles in the helical assembly, which will be described in detail below.

As show in Fig. 1c and 1d, the first and second layers at one end of the IL-17RA/RB/ACT1 helix is formed by the SEFIR domains of IL-17RB and IL-17RA, respectively, while the remaining layers are all occupied by ACT1. Placing this end of the helix below the plasma membrane allows the SEFIR domain of IL-17RB to connect to the transmembrane regions through the ∼17-residue flexible linkers between them, based on the AlphaFold model of full-length IL-17RB (Fig. 1d)^24,25^. The IL-17RA SEFIR domain in the second layer is further away from the membrane (Fig. 1d). Accordingly, the linker between the transmembrane and SEFIR domains in IL-17RA is longer (∼31 residues) and sufficient to span the distance between the two domains.

### The hetero-tetramer of the SEFIR domains of IL-17RA and IL-17RB

The SEFIR domain of IL-17RB in the IL-17RA/RB/ACT1 complex is similar to the previously published crystal structure, adopting the typical SEFIR/TIR domain fold with five β-strand-loop-ɑ-helix modules organized into a globular fold (Supplementary Fig. 4b)^21–23,26^. We follow the same nomenclature used in those studies to describe the SEFIR domain structures of IL-17RA, RB and ACT1, where the five β-strands (βA-E) are surrounded by the five ɑ-helices (ɑA-E) (Supplementary Fig. 4b). The cryo-EM structure suggests that C343 and H346 from ɑA and C404 and C408 from the loop between βC and ɑC (the CC loop) together form a Zn^2+^ binding site, which is disordered in the crystal structure (Supplementary Fig. 4c). This Zn^2+^ binding site does not participate in the protein-protein interactions in the complex but may play a role in stabilizing the structure.

The cryo-EM structure shows that the SEFIR domain of IL-17RA has an extra β-strand-ɑ-helix module, βF and ɑF, compared to IL-17RB (Supplementary Fig. 4b, 4e). βF and the preceding EF loop together form a hairpin (referred to as the β-hairpin thereof) in the IL17-RA/RB/ACT1 complex, which plays an important role in recruiting ACT1 by forming a small β-sheet with the C-terminal extension of ACT1 from the next layer in the filament (See detail below). AlphaFold correctly predicts the β-hairpin of IL-17RA, which closely resembles that in the cryo-EM structure (Supplementary Fig. 4e). However, in the previously published crystal structure of IL-17RA SEFIR domain, while ɑF is also present, βF and the preceding EF loop are disordered (Supplementary Fig. 4c)^27^. IL-17RA therefore likely has propensity to form the β-hairpin, although it is unstable without the interactions with IL-17RB. Experimental structures of the SEFIR domains of IL-17RC, RD and RE are not available. AlphaFold predicted models of these IL-17Rs show that they all have ɑF, but the preceding sequence does not form a β-hairpin (Supplementary Fig. 4e). These observations together suggest that the β-hairpin is one of the unique structural elements in IL-17RA that enables it to act as the ACT1-binding component in heteromeric IL-17R complexes.

The hetero-tetramer of IL-17RA/RB is mediated by multiple inter-domain interfaces similar to those seen in the TIR domain filaments (Fig. 2a and Supplementary Fig. 4a)^21–23^. The inter-strand interface between the two IL-17RB molecules is formed by helices ɑB and ɑC (BC surface) from the top IL-17RB molecule and helices ɑC and ɑD (CD surface) from the second (referred to as IL-17RB’) (Fig. 2b). V379, A383, L425 and L429 from IL-17RB and F427, C431, L434, A459 and

**Fig. 2.**
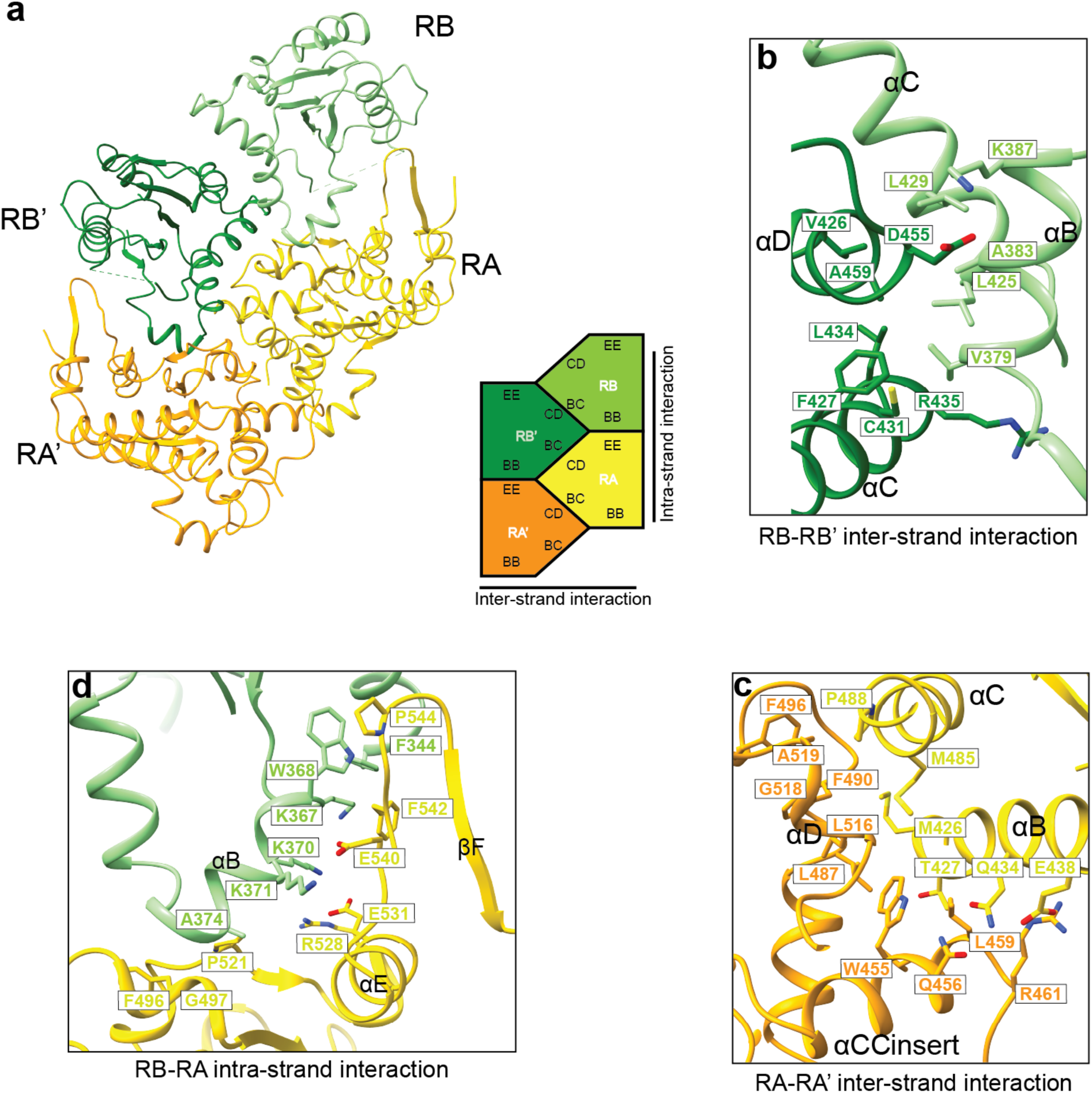
IL-17RA/RB SEFIR domain hetero-tetramer in the IL-17RA/RB/ACT1 complex. **a**, Overall structure of the hetero-tetramer in the cartoon and schematic representations. **b**, Detailed view of the inter-strand interaction between the two IL-17RB SEFIR domains. **c**, Detailed view of the inter-strand interaction between the two IL-17RA SEFIR domains. **d**, Detailed view of the intra-strand interaction between the SEFIR domains of IL-17RB and IL-17RA.

V462 from IL-17RB’ together constitute the hydrophobic core of the interface. Similarly, the inter-strand interface in the second layer is formed by the BC surface of IL-17RA and the CD surface of IL-17RA’ (Fig. 2c). The core of this interface contains M426, T427, M485 and P488 from IL-17RA and W455, L459, L487, F490, F496, L516, G518 and A519 from IL-17RA’. Both W455 and L459 are located in the long loop between βC and ɑC, which forms a 3-turn helix that is not a typical feature of the SEFIR/TIR fold (ɑCCinsert in Fig. 2c). Both the inter-strand interfaces of IL-17RB/IL-17RB’ and IL-17RA/IL-17RA’ are augmented by numerous polar interactions at the periphery. In addition, IL-17RB’ and IL-17RA form an inter-strand interface, between the BC face of IL-17RB’ and the CD face of IL-17RA. The binding mode and residues involved are similar to those of the IL-17RB/IL-17RB’ and IL-17RA/IL-17RA’ inter-strand interfaces.

The intra-strand interaction in the IL-17RA/RB hetero-tetramer is mostly mediated by the BB face of IL-17RB and the EE face of IL-17RA (Fig. 2d). The last and first ordered residues from IL-17RB and IL-17RA respectively is ∼33 Å apart, which could be readily spanned by the ∼30 disordered linker residues in the IL-17RB-RA fusion protein. Therefore, the linker facilitates the interaction between IL-17RA and IL-17RB by increasing their local concentrations but does not impose artificial restraints. In the interface, the long BB-loop from IL-17RB forms a 2-turn helix, which docks on the relatively flat EE surface of IL-17RA through both hydrophobic and electrostatic interactions (Fig. 2d). This binding mode places the β-hairpin in IL-17RA in proximity with the beginning portion of the BB-loop in IL-17RB. W368 in this loop of IL-17RB makes a stacking interaction with P544 in the IL-17RA β-hairpin. A cation-π interaction is formed between K367 from IL-17RB and F542 in the β-hairpin. Two ion-pairs between, IL-17RB^K371^–IL-17RA^E540^ and IL-17RB^K370^–IL-17RA^E531^, further contribute to the interaction (Figure 2d). These interactions stabilize the β-hairpin, allowing it to mediate the interaction with ACT1 (See below for details). Interestingly, W368 is conserved in other IL-17Rs, except for IL-17RA which has a leucine at the equivalent position (Leu415) (Supplementary Fig. 4d). This tryptophan residue appears to be a signature feature in the co-receptors (IL-17RB, RC, RD and RE) that stabilizes the β-hairpin in IL-17RA for recruiting ACT1.

### Interactions between IL-17RA and ACT1

The formation of the IL-17RB/RA hetero-tetramer promotes the recruitment of ACT1 through several mechanisms. Firstly, it creates a docking site where ACT1 SEFIR domain can simultaneously engage with the two IL-17RA SEFIR domains (Fig. 3a). The intra-strand interaction is formed between the BB-face of IL-17RA and the EE-face of ACT1. This interface contains a mixture of hydrophobic and polar residues, including S421, Q418, Pro474 from IL-17RA, and M512, F514, N535, T536 and H537 from ACT1 (Fig. 3b).

**Fig. 3.**
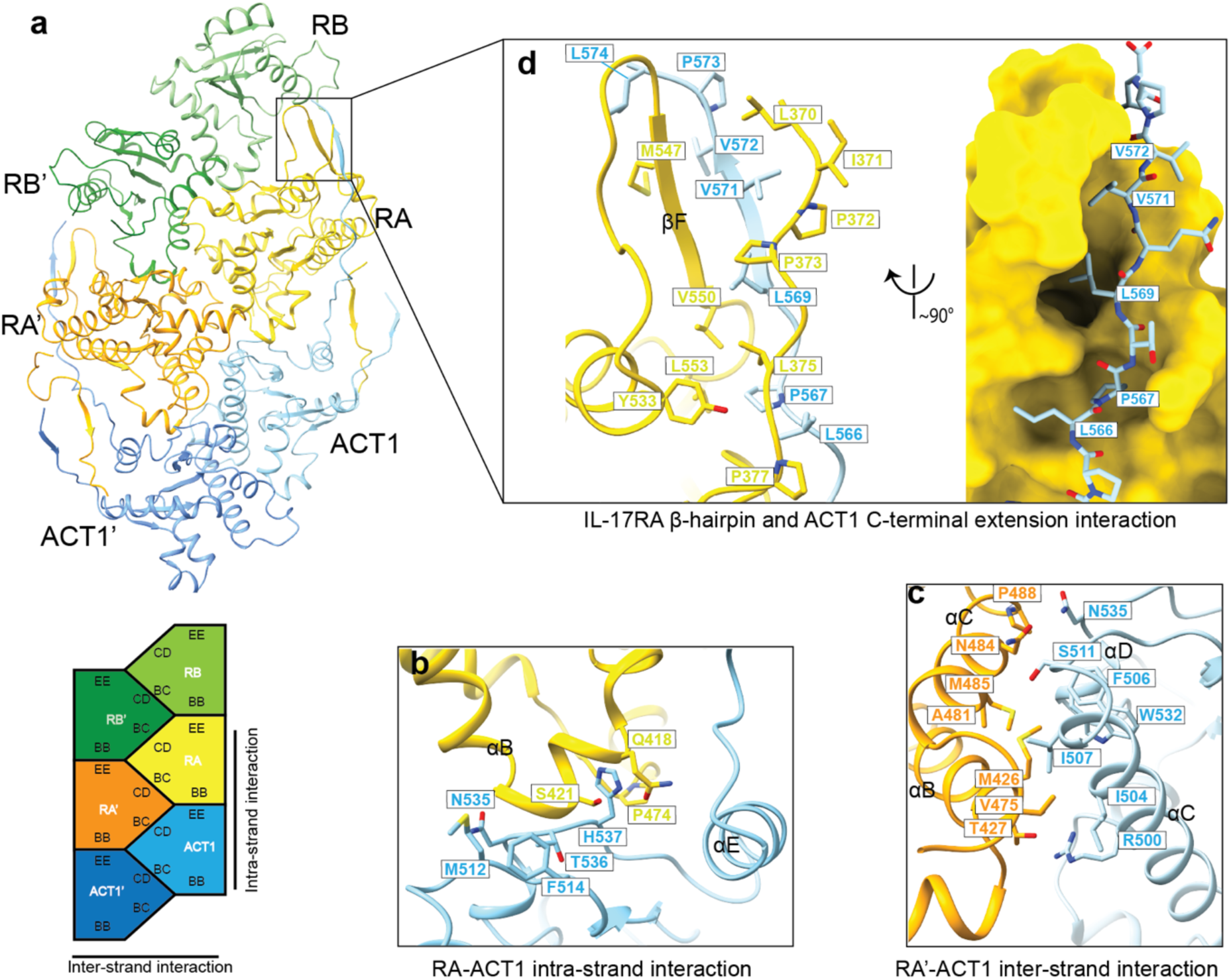
Interactions between the IL-17RA SEFIR domain and ACT1. **a**, Overall structure of the IL-17RA/RB/ACT1 complex in cartoon and schematic representations. For simplicity, only two ACT1 protomers are shown to illustrate their interactions with IL-17RA. **b**, Detailed view of the intra-strand interaction between the SEFIR domains of IL-17RA and ACT1. **c**, Detailed view of the inter-strand interaction between the SEFIR domains of IL-17RA and ACT1. **d**, Detailed view of the interaction between the β-hairpin of IL-17RA and the C-terminal extension of ACT1.

Notably, a biallelic T536I mutation of ACT1 has been found to cause chronic mucocutaneous candidiasis in human patients^28^. Based on the structure, this mutation likely drives the disease by destabilizing this IL-17RA/ACT1 interface and thereby impairing the IL-17R immune signaling. The inter-strand interaction between the BC surface of IL-17RA’ and the CD face of ACT1 is more hydrophobic at its core, containing residues M426, V475, A481, M485 and P488 from IL-17RA’ and I504, F506, I507, W532 from ACT1 (Fig. 3c). Several polar residues from both IL-17RA and ACT1 also contribute to this interface.

The second major mechanism that the IL-17RA/RB hetero-tetramer uses to enhance ACT1 recruitment is the stabilization of the β-hairpin in IL-17RA by IL-17RB. The C-terminal extension of the ACT1 SEFIR domain in the first ACT1 layer extends across the SEFIR domain surface of IL-17RA (Fig. 3d). The last 6 residues (569-574) in ACT1 adopts a β-strand conformation and forms a small β-sheet with the β-hairpin in IL-17RA. In addition, the N-terminal extension of IL-17RA (residues 370-377), the β-hairpin and surrounding residues in IL-17RA together form a hydrophobic groove, which binds the hydrophobic residues in the ACT1 C-terminal extension (Leu566, Pro567, Leu569, Val571 and Val572) (Fig. 3d). This tethering interaction between the C-terminal extension of ACT1 and the β-hairpin of IL-17RA is unique to IL-17RA/ACT1, with no similar interactions observed in the TIR domain oligomers.

### Stabilization of the SEFIR-SEFIR interaction of ACT1 by the SEFEX of IL-17RA

The interactions among the ACT1 SEFIR domains in the helical assembly are similar to those among IL-17RB, RA and ACT1 described above (Fig. 4). The intra-strand interface involves D445, G449, I450 and D451 in the BB-surface of one ACT1 SEFIR and M512, N513, F514, N535, T536, H537 in the EE-surface of the second ACT1 SEFIR (ACT1’) (Fig. 4b). The inter-strand interface is mediated by D451, I453, M456, E457, L493, Y497, M501 and I504 in the BC-face of the first ACT1 and R500, Q503, I504, F506, I507, S511, G510 and T531 in the CD-face of the second ACT1 (ACT1^2^) (Fig. 4c).

**Fig. 4.**
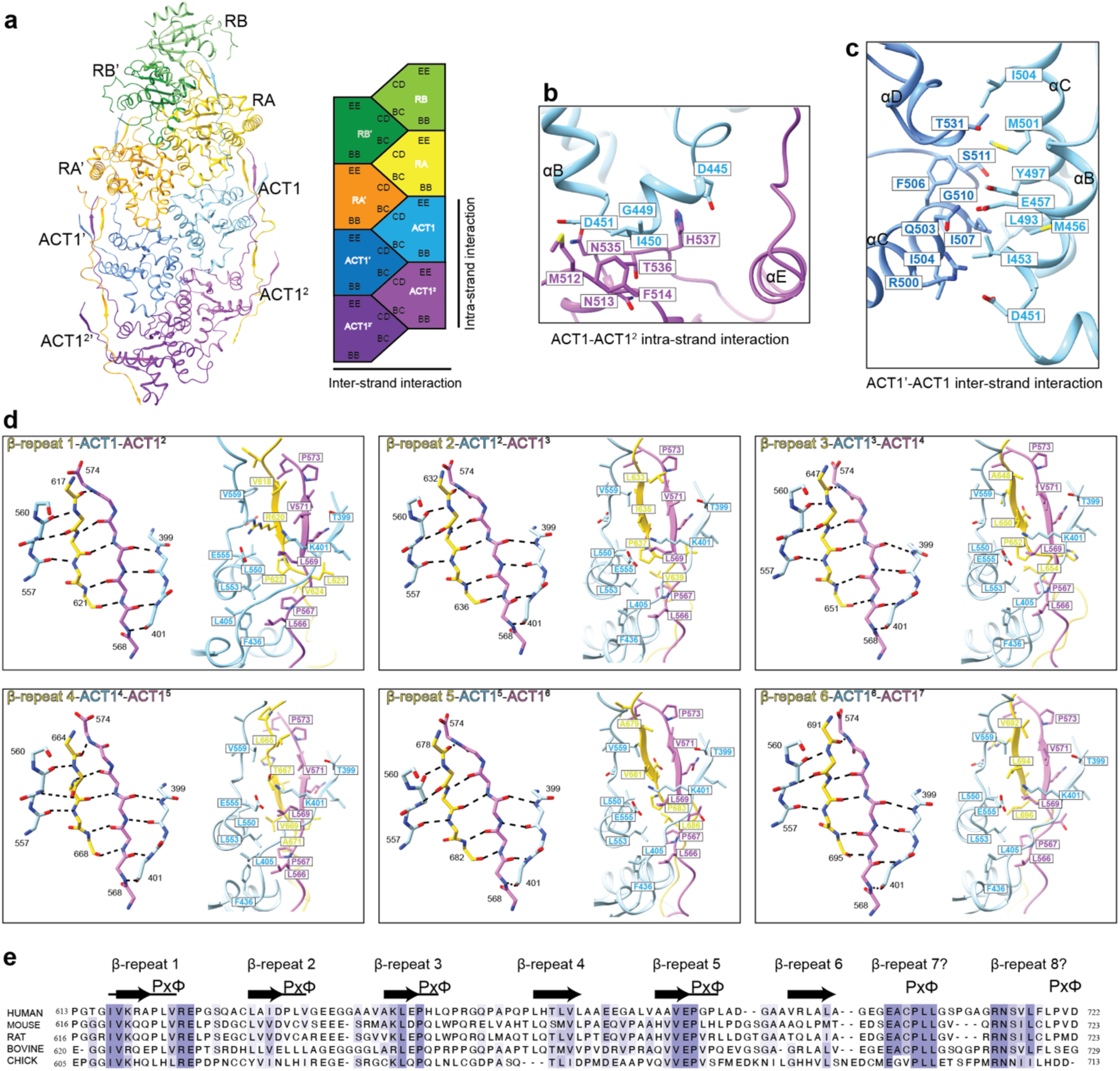
Interactions between ACT1 protomers and between ACT1 and the IL-17RA SEFEX. **a**, Overall structure of the IL-17RA/RB/ACT1 complex in cartoon and schematic representations. For simplicity, only four ACT1 protomers are shown. **b**, Detailed view of the intra-strand interaction between ACT1 protomers. **c**, Detailed view of the inter-strand interaction between ACT1 protomers. **d**, Detailed views of the interactions between the β-repeats in the SEFEX of IL-71RA and ACT1 protomers in the IL-17RA/RB/ACT1 filament as shown in Fig. 1d. **e**, Sequence alignment of the SEFEX of IL-17RA from different species. The degenerate β-strand-PxΦ repeats are highlighted. Question marks indicate that the sequence conservation pattern suggests the presence of β-repeats 7 and 8, but they are not resolved in the cryo-EM structure.

In addition to the SEFIR-SEFIR interactions, the N- and C-terminal extensions of the ACT1 SEFIR and the SEFEX of IL-17RA together form a small anti-parallel β-sheet beside each ACT1 to stabilize the helical assembly. For the first ACT1 molecule below IL-17RA, the β-sheet is formed by the following four structural elements, in the order of increasing distance from the SEFIR domain: (1) the beginning portion (residues 557-560) of the C-terminal extension of the current ACT1, (2) the end portion (residues 568-574) of the C-terminal extension from the ACT1 molecule from the layer below (ACT1^2^), (3) residues 617-621 from the middle of IL-17RA SEFEX, and (4) residues 399-401 from the N-terminal extension of the current ACT1 SEFIR (Fig. 4d). Residues in the β-sheet make both backbone hydrogen bonds and sidechain packing and electrostatic interactions. In addition, the middle portion (residues 561-568) of the C-extension of ACT1^2^ makes numerous interactions with the body of the current ACT1 SEFIR domain. Similarly, the segment (residues 622-631) in the IL-17RA SEFEX following the β-strand portion, including a ^622^PLV^624^ motif, wraps around the current ACT1 SEFIR domain and make numerous contacts.

The atomic model contains the β-sheet module for six ACT1 molecules in each strand, whereas this feature is absent or only partially modelled for the three ACT1 subunits located at the distal end of the helical assembly due to weak density (Fig. 1c and 4e). The modules by each ACT1 are similar, except that they contain different segments of the SEFEX of IL-17RA. The structure therefore reveals that the SEFEX of IL-17RA contains at least 6 hidden repeats that can form the β-sheet module, which are spaced ∼15-residue apart. A sequence alignment of the IL-17RA from different species shows that the β-strand forming portion of the repeats are more conserved than the flanking spacers (Fig. 4e). Notably, the repeats also contain a degenerate PxΦ motif (x, any residue; Φ, hydrophobic) following the β-strand, corresponding to ^622^PLV^624^ in the first repeat mentioned above. Additional repeats beyond repeat 6 may exist in the SEFEX based on the sequence alignment and the presence of longer filaments in the micrographs, but they were not modelled due to poor cryo-EM density at the distal end of the helical assembly (Fig. 4e).

### Mutational analyses of the IL-17RA/RB/ACT1 helical assembly

The cryo-EM structure of the IL-17RA/RB/ACT1 helical assembly shows how the receptor hetero-tetramer organizes the two IL-17RA molecules to promote the oligomerization of ACT1. To validate the structural model, we designed mutations in IL-17RA, IL-17RB and ACT1 at each of the binding interfaces and tested their effects on the formation helical assembly. For IL-17RB, we mutated W368, which interacts with the β-hairpin in IL-17RA, to alanine, glutamate and leucine respectively. In addition, a V379R mutation was introduced at the IL-17RB inter-strand interface. Mutations of ACT1 included I507R (RA-ACT1 and ACT1-ACT1 inter-strand interface) and V559P, L566E/P567E, Δ570-574 (ΔC) in the C-terminal extension. We also made the disease-associated T536I mutation in ACT1 as described above. Finally, mutations of IL-17RA included G497L (RB-RA intra-strand interface), S421R (RA-ACT1 intra-strand interface), M426E, M485E, P488E (RA-RA, RA-ACT1 inter-strand interface), F542R/P544A (β-hairpin), P622E (β-repeat 1 in the SEFEX). The mutations were individually introduced into the IL-17RB-RA fusion or ACT1 constructs for co-expression in HEK293F cells as for the wild type described above. The results showed that each of these mutations abolished the formation of the IL-17RA/RB/ACT1 complex (Supplementary Fig. 5). Consistently, the protein samples containing any of the mutations failed to form filaments as shown by negative-stain EM (Fig. 5). These results demonstrate that the binding interfaces seen in the cryo-EM structure are required for the formation of the IL-17RA/RB/ACT1 helical assembly.

**Fig. 5.**
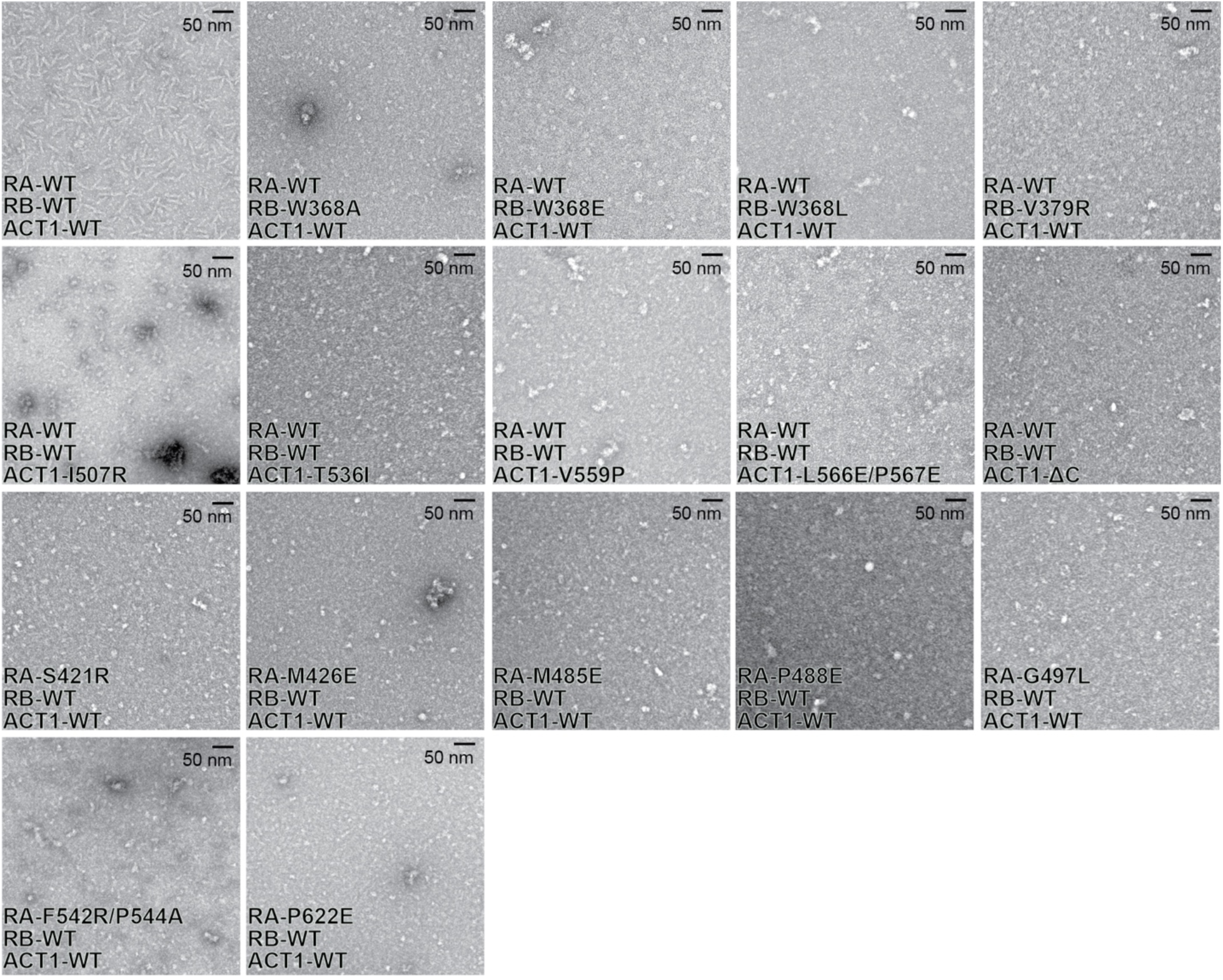
Negative-stain EM images showing that mutations in IL-17RA, RB and ACT1 abolish the formation of helical assembly. For the sample of wild type IL-17RA, RB and ACT1, the protein complex in fractions 2-4 in Supplementary Fig. 5a were imaged. For the samples with mutated proteins, fractions 2-4 contained little protein, images were collected on pooled fractions 5-10 instead.

To test the effects of the mutations on the formation of the IL-17RA/RB/ACT1 complex in cells, we carried out co-immunoprecipitation experiments using HEK293T cells co-transfected with epitope tagged full-length IL-17RA, RB and ACT1. As shown by previous studies, ACT1 could co-immunoprecipitate with the IL-17 receptors from these cells without IL-17 treatment, likely due to constitutive low levels of secretion of the IL-17 ligands by these cells^16,20^. Our results showed that both IL-17RA and ACT1 robustly co-immunoprecipitated with IL-17RB (Supplementary Fig. 6a). The amounts of ACT1 pulled down by IL-17RB were reduced substantially by the interface mutations in IL-17RA (F542R/P544A and P622E/P637E/P652E), IL-17RB (W368A) or ACT1 (T536I and ΔC). The amount of IL-17RA pulled down appeared not affected by any of the mutations, presumably because the interaction of the extracellular domains of IL-17RA and RB induced by the ligand were sufficiently strong to support their co-immunoprecipitation^13^.

### The IL-17RA/RB/ACT1 helical assembly is important for signaling in cells

To examine whether the helical assembly underlies the activation and signaling of ACT1 in cells, we adopted a previously established cell-based assay, which assesses IL-17-stimulated IL-6 secretion in HeLa cells^17,29^. Cells co-expressing IL-17RA, RB and ACT1 through transient transfection showed robust increase in IL-6 secretion in response to IL-25 treatment, whereas IL-6 production from cells un-transfected or transfected with only one of the three proteins was much lower (Supplementary Fig. 7). The four mutations of IL-17RB described above reduced the IL-6 production to near the basal level from cells transfected with IL-17RA and ACT1 but not IL-17RB (Fig. 6a). The mutations of ACT1 had similar negative effects on IL-6 production (Fig. 6b). For testing effects of IL-17RA mutants, we used siRNA to knock down the endogenously expressed IL-7RA (Supplementary Fig. 6c). The results showed that all tested single mutations except S421R significantly diminished IL-6 secretion, although the reductions were not as pronounced as the mutations in IL-17RB or ACT1 (Fig. 6c). We further tested several double and triple mutations of IL-17RA, and a mutant containing a deletion of residue 611-866 in the SEFEX region (ΔSEFEX). These mutants exhibited more pronounced decrease in IL-6 secretion compared to the single mutants. As a control, we show that all the mutants of IL-17RA and RB were expressed at levels comparable to their respective wild-type counterparts, while the mutants of ACT1 expressed at higher levels than the wild type (Supplementary Fig. 6b). Therefore, the mutations caused reduced IL-6 secretion by disruption of the IL-17RA/RB/ACT1 helical assembly, rather than destabilization or reduced expression of the proteins.

**Fig. 6.**
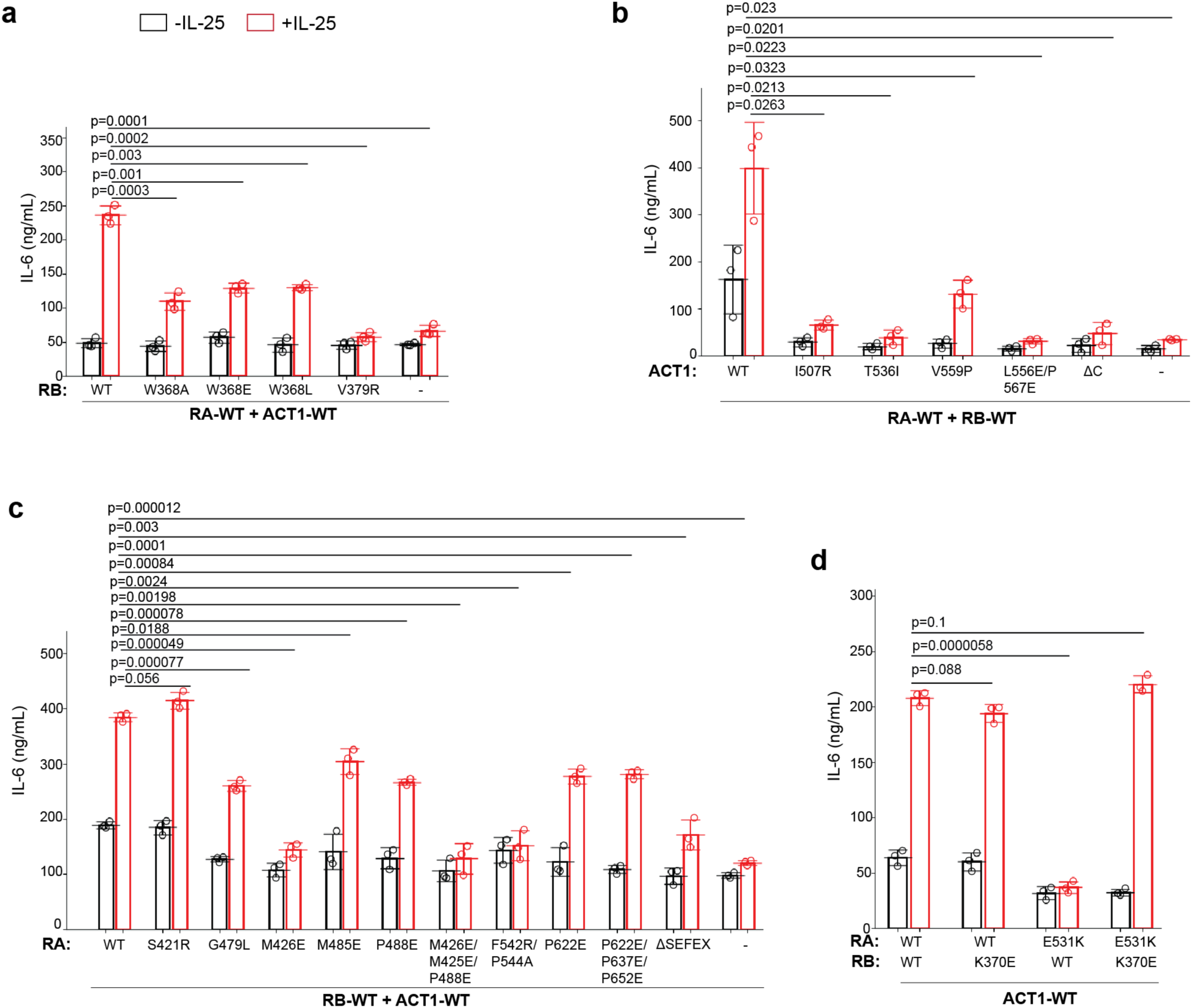
Mutational analyses of the IL-17RA/RB/ACT1 helical assembly in IL-25-stimulated IL-6 secretion in HeLa cells. **a,** Effects of mutations in IL-17RB on IL-6 production. **b,** Effects of mutations in ACT1 on IL-6 production. **c,** Effects of mutations in IL-17RA on IL-6 production. **d,** Rescue of IL-6 production by a pair of charge-swapping mutations in IL-17RA and IL-17RB. For **a** and **b,** cells were co-transfected with the indicated combinations of the IL-17RA, IL-17RB and ACT1 constructs. For **c** and **d**, cells were first transfected with siRNA-1 (Supplementary Fig. 6c) to knock down endogenous IL-17RA expression and then transfected with indicated plasmids. Cells were stimulated with IL-25 at 25 nM. IL-6 levels in the medium were measured with ELISA. Each circle represents the result from one of three independent experiments. The bars represent means and standard deviations. P-values were calculated with two-tailed Student’s T-test.

In addition, we designed a pair of charge-swapping mutations to further test the binding mode between IL-17RA and RB. As mentioned above, K370 in IL-17RB and E531 in IL-17RA form an intra-strand charge-charge interaction (Fig. 2d). E531 also makes intra-molecule contacts with R528 in IL-17RA. As a results, this part of the IL-17RA/RB interface contains two positively charged residues and one negatively charged residue. The K370E mutation of IL-17RB did not have a significant effect on IL-6 secretion (Fig. 6d), suggesting that the IL-17RA/RB complex could tolerate the loss of the charge-charge interaction between these two residues. In contrast, the E531K mutation of IL-17RA completely abolished IL-25-induced IL-6 production in HeLa cells (Fig. 6d). With the E531K mutation, this part of the interface contains three positively charged residues (K370 in IL-17RB, K531K and R528 in IL-17RA), leading to strong electrostatic repulsion and destabilization of the IL-17RA/RB complex, which likely underlies the abolished IL-6 secretion. Consistent with this interpretation, co-expression of wild type ACT1 with the IL-17RA E531K and IL-17RB K370E mutants, which restored the charge ratio at the IL-RA/RB interface, rescued IL-25-stimulated IL-6 secretion to the wild type levels (Fig. 6d). Taken together, these results demonstrate that the helical assembly of IL-17RA/RB/ACT1 is the basis for the activation of ACT1 by IL-17R hetero-tetramers in cells.

### IL-17RA, RC and ACT1 form the same type of helical assembly

Interestingly, an AlphaFold predicted complex of two IL-17RA, two RC and 12 ACT1 molecules is very similar to our cryo-EM structure of the IL-17RA/RB/ACT1 complex (Fig. 7a), suggesting that IL-17RC uses the same mechanism to form the complex with IL-17RA and activate ACT1. Consistent with this model, the co-expressed IL-17RC-RA fusion and ACT1 formed a large oligomeric complex in gel filtration chromatography (Fig. 7b and 7c). Negative-stain images showed that filaments of the IL-17RA/RC/ACT1 complex were similar to those of IL-17RA/RB/ACT1 (Fig. 7d). We further tested this model by using the IL-6 production assay in HeLa cells. Cells transfected with wild type IL-17RA, RC and ACT1 showed robust IL-6 production in response to IL-17F stimulation (Fig. 7e). Mutations of W549L in IL-17RC, corresponding to W368L in IL-17RB, led to significant reduction of IL-6 secretion (Fig. 7e). Similarly, the mutation of V560R in IL-17RC, corresponding to V379R in IL-17RB, also significantly reduced IL-6 production (Fig. 7e). These results strongly suggest that IL-17RA, RC and ACT1 also form the left-handed double-helix assembly similar to that of IL-17RA, RB and ACT1.

**Fig. 7.**
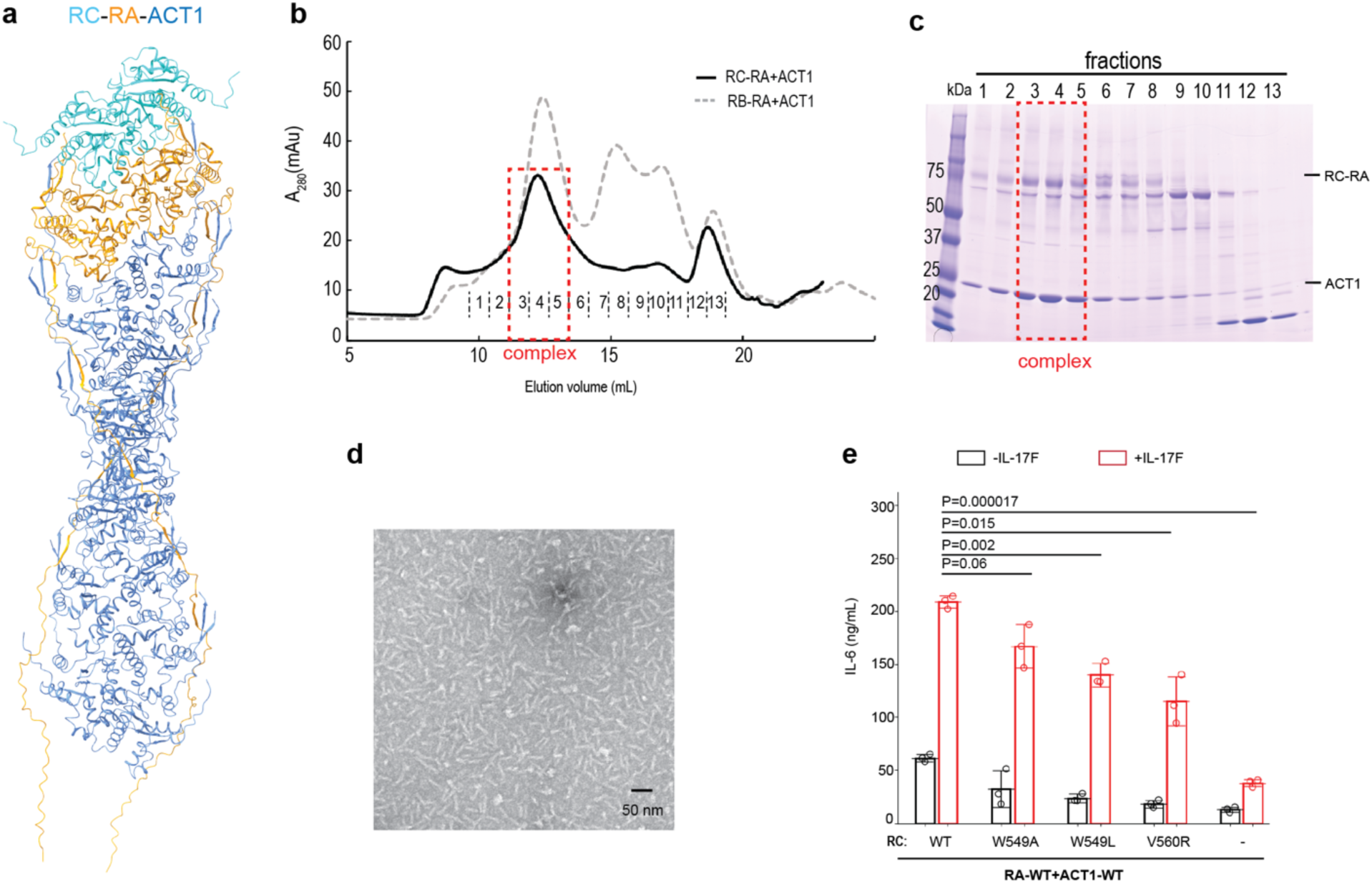
Formation of the helical assembly by IL-17RA, RC and ACT1. **a**, AlphaFold predicted model of the IL-17RA/RC/ACT1 complex. **b**, Gel filtration chromatography profile of the IL-17RA/RC/ACT1 complex. **c**, SDS-PAGE analyses of fractions from **b**. The result is a representative of three repeats. **d**, Negative-stain image of the IL-17RA/RC/ACT1 complex. The result is a representative of three repeats. **e**, IL-6 production assay of HeLa cells transfected with different combinations of IL-17RA, RC and ACT1. The experiments were carried out in the same manner as in Fig. 6. Each circle represents the result from one of three independent experiments. The bars represent means and standard deviations. P-values were calculated with two-tailed Student’s T-test.

## Discussion

Formation of large signalosomes through helical assemblies of small globular domains such as DEATH, TIR and SEFIR domains is a reoccurring theme in innate immunity^21–23,30,31^. In this study we present the cryo-EM structure of the IL-17RA/RB/ACT1 complex, revealing both the similarities and differences between this SEFIR domain-mediated helical assembly and the related TIR domain oligomers^21–23^. The IL-17RA/RB/ACT1 helical assembly described here provides a structural basis for why the oligomerization and activation of ACT1 requires the hetero-tetramerization between IL-17RA and one of the co-receptors^9,10,12,13^. IL-17RB (and likely other co-receptors) does not directly interact with ACT1 but contributes to the helical assembly by priming the two IL-17RA molecules for recruiting ACT1. The conserved tryptophan residue (W368 in RB) in the co-receptors is key to their function, by stabilizing the β-hairpin in IL-17RA for binding

ACT1. The structure reveals that at least six degenerate β-PXΦ motifs in the SEFEX region of IL-17RA form repetitive β-sheet modules with both the N- and C-terminal extension of ACT1. This remarkable tendril-like function of the IL-17RA SEFEX in binding ACT1 and stabilizing the helical assembly provides an explanation of its important role in signaling^20^. The helical assemblies formed by IL-17Rs and ACT1 appear stable and may not readily dissociate to terminate the signal. Previous studies have shown that termination of the IL-17R/ACT1 signal could be mediated by other mechanisms such as ubiquitination and degradation of ACT1, and phosphorylation of ACT1 that prevents binding of downstream transducers^19,32^.

To address the question whether other IL-17R family co-receptors, RD and RE, form the same helical assembly with IL-17RA and ACT1, we carried out AlphaFold predictions of the complexes of IL-17RA, the coreceptors and ACT1. The predicted complexes were both similar to the cryo-EM structure (Supplementary Fig. 8), suggesting that the helical assembly is the conserved mechanism for the activation of ACT1 by different combinations of IL-17RA and the co-receptors. It however appeared that successful prediction was dependent on inputs with the correct stoichiometry of IL-17RA, the co-receptor and ACT1 based on our cryo-EM structure. For example, AlphaFold failed to predict the individual homo- or hetero-dimeric interactions among these proteins. Therefore, experimental data were critical for understanding the configuration of the complexes in this case, while AlphaFold provided valuable additional insights. The C-terminal extensions in the co-receptors, except IL-17RD, are short (fewer than 30 residues) and unlikely possess the ability of interacting with ACT1 as the IL-17RA SEFEX. However, the C-terminal extension in IL-17RD is ∼170-residue long, comparable to the IL-17RA SEFEX. Interestingly, AlphaFold predicted model of 2 copies of the IL-17RD SEFIR with the extension region and 8 copies of ACT1 shows an assembly similar to the IL-17RA/RB/ACT1 cryo-EM structure, in which the C-terminal tail of IL-17RD makes two β-sheet modules with ACT1 (Supplementary Fig. 8 and 9). These analyses raise the question whether IL-17RD may directly interact with ACT1, particularly considering that IL-17RD is the most ancient IL-17R that might have functioned independently in activating ACT1 before the emergence of the specialized common receptor IL-17RA and co-receptors in the IL-17R family^8^.

It is intriguing that various combinations of IL-17 and IL-17Rs form heteromeric assemblies with completely different arrangements of the receptor ectodomains^12,13^, seemly conflicting with the idea of a unified mechanism for inducing ACT1 oligomerization. Our results show that increasing local concentration of the IL-17RA and RB SEFIR domains by a flexible linker between the two is sufficient to promote the hetero-tetramer formation and the oligomerization of ACT1. Taken together, it seems that the IL-17 ligand activates the signaling pathway by increasing the local concentration of IL-17RA and a co-receptor and promoting their hetero-tetramerization or high-order oligomerization, but it does not dictate the specific binding mode among the intracellular SEFIR domains of the receptors. The flexible linkers flanking the transmembrane region of the receptors allow the intracellular SEFIR domains to self-organize and form the helical assembly with ACT1 through the intrinsic specific interactions among themselves. High-order oligomers of ACT1 recruit multiple TRAF6 to form large signalosomes, which efficiently activates NFκB and promotes inflammatory signaling^31^.

The IL-17 pathway has been extensively targeted for treating inflammatory diseases such as psoriasis^4,5^. Current treatments primarily rely on antibodies against IL-17 or IL-17Rs, which are effective but expensive and require injection. The structure presented in this study provides a foundation for future development of small-molecule drugs that inhibit IL-17 signaling by disrupting the double-stranded helical assembly of the IL-17RA/co-receptor/ACT1 complexes.

## Methods

### Protein expression and purification

The cDNAs encoding *human* IL-17RA (Uniprot code: Q96F46-1), IL-17RB (Uniprot code: Q9NRM6-1), Act1 (Uniprot code: O43734-1) and IL-25 (Uniprot code: Q9H293-1) were purchased from The McDermott Center for Human Growth and Development in UT Southwestern Medical Center. The DNA encoding *human* IL-17RC (Uniprot Code: Q8NAC3-2) was synthesized by IDT. The IL-17RA-RB fusion construct in the pEZT-BM vector contained the coding regions for the T6SS immunity protein 3 (Tsi3) from Pseudomonas aeruginosa as a purification tag^33,34^, the cleavage site for the *human* rhinovirus 3C protease, residues 328-484 of IL-17RB, a 4x(GGGS) flexible linker and finally residues 361-866 of IL-17RA. The RC-RB fusion had a similar architecture, except that residues 510-705 of IL-17RC replaced the segment of IL-17RB and the linker was 3x(GGGS). The coding region for residues 372-574 of ACT1 was cloned into the same vector. Mutations were introduced by polymerase chain reaction-based mutagenesis. A previous study showed that Y558 in ACT1 could undergo phosphorylation^29^. To avoid potential heterogeneity as a result of this phosphorylation, the Y558F mutant of ACT1 was used for the cryo-EM sample. Other experiments including the negative-staining EM and the functional assays used ACT1 constructs that did not contain this mutation.

ACT1 was co-expressed with the wild type or mutants of IL-17RB-RA or IL-17RC-RA fusions in HEK293F cells cultured in suspension in FreeStyle293 expression medium (Gibco, 12338-018) through transient transfection. The two plasmids (500 µg each) were used with 3 ml polyethylenimine (PEI) at 1 mg/ml for transfecting 1 L cells. These and other cells used were assumed to be authenticated by the commercial sources, therefore were not further authenticated in the study. Cells were collected 72 h after transfection by centrifugation, resuspended in buffer A (50 mM Tris-HCl pH 8, 150 mM NaCl, 5% glycerol, 1% DNase I, 0.2 mM AEBSF, 2 mM ϕ3-ME), and lysed by French press. Lysates were centrifuged to remove debris. The affinity purification step of IL-17RB-RA and ACT1 was based on the high-affinity interaction between the N-terminal Tsi3 tag and the T6SS effector protein Tse3^34^. The target proteins were captured by Tse3-conjugated Sepharose 4B resin (GE Healthcare) equilibrated in buffer B (50 mM Tris-Cl pH 8, 150 mM NaCl, 5% glycerol, 2 mM ϕ3-ME). Unbound proteins were removed by extensive wash with buffer B. Target protein was eluted by cleavage from the Tsi3 tag with the 3C protease on resin at 4 °C for 12 h. The elute was further purified on a Superose 6 Increase 10/300 GL (GE Healthcare) in buffer C (20 mM HEPES pH 7.5, 150 mM NaCl, 1 mM DTT). Peak fractions were pooled, concentrated and kept at −80 °C before use. For expression of IL-25 (IL-17E), the coding region for residues 30-177 of IL-25 was cloned into the pEZT-SP vector, which is the same as pEZT-BM but encodes a signal peptide at the N terminus and an 8xHis tag at the C terminus. The plasmid was transfected into HEK293F cells using PEI for protein expression. Six days after transfection, the medium was collected and incubated with Ni Sepharose™ Excel beads (Cytiva). The purification procedure for IL-25 was similar to that described previously^35^. Briefly, protein captured by the Ni beads was eluted with a buffer containing 20 mM Tris-HCl pH 8.0, 500 mM NaCl and 500 mM imidazole. The eluted protein was further purified on a Superdex 75 10/300 column (GE Healthcare) in a buffer containing 20 mM HEPES pH 7.5 and 150 mM NaCl. Peak fractions were pooled, concentrated and kept at −80 °C before use.

### siRNA Knock-down of IL-17RA

HeLa cells were cultured in Dulbecco’s modified Eagle’s medium (DMEM; Gibco) supplemented with 10% fetal bovine serum (FBS; Gibco) and 1% penicillin–streptomycin at 37°C in 5% CO₂. Cells were seeded in 12-well plates to reach ∼40% confluence at the time of transfection. Two pairs of small interfering RNA (siRNA) targeting 3’-UTR of IL17RA were tested. The sequences of the first (siRNA-1) were AGGAGUGUGUGUGCACGUAdTdT (sense) and UACGUGCACACACACUCCUdTdT (anti-sense, while the sequences of the second (siRNA-2) were GUGUAUAUGUUCGUGUGUGdTdT (sense) and CACACACGAACAUAUACACdTdT (anti-sense). The non-targeting control had the sequences of GUGUGAUGAUCCUGAAGCUdTdT (sense) and AGCUUCAGGAUCAUCACACdTdT (anti-sense). Cells were transfected with the siRNA using Lipofectamine 3000 (Invitrogen). Forty-eight hours after transfection, cells were lysed in the RIPA buffer (Cat# 20-188, Sigma). Proteins were extracted and IL-17RA was detected by western blot using anti-IL-17RA antibody (Cat# 12661T, cell signaling) and the secondary antibody anti-rabbit IgG (H+L) (DyLight 800 4X PEG Conjugate) (Cat#5151S, cell signaling). The results showed that both the IL-17RA siRNAs successfully knocked down the expression of IL-17RA (Supplementary fig. 6c). siRNA-1 was used for knocking down IL-17RA in the following experiments.

### IL-6 production assay in HeLa cells

Full-length IL-17RA and IL-17RC were cloned into the pETZ vector with a C-terminal His_8_-PA tag^36^. Full-length IL-17RB was cloned into the pEZT vector with a C-terminal His_8_-Tsi3 tag. Full-length ACT1 was cloned into the pEZT vector with a N-terminal His_8_-PA tag. Mutations were introduced by PCR-based mutagenesis. Hela cells were grown in 12-well plates to ∼70–80% confluency before transfection. For assays with IL-25 stimulation, cells were co-transfected with the plasmids of IL-17RA, IL-17RB and ACT1 (10 ng each) and 220 ng empty vector using Lipofectamine 3000 (Invitrogen). For testing the IL-17RA mutants, cells were first transfected with siRNA-1 to knock down the endogenous IL-17RA expression (see above). One day after siRNA transfection, cells were co-transfected with the IL-17RA, RB and ACT1 plasmids. For assays with IL-17F stimulation, the IL-17RC plasmids were used in the place of IL-17RB. Culture medium was replaced with fresh DMEM (Thermo Fisher Scientific) 6 hours after transfection. Cells were treated with IL-25 or IL-17F at 25 nM after the medium exchange. Eighteen hours later, medium was collected for Enzyme-Linked ImmunoSorbent Assay ELISA assays for IL-6 using Human IL-6 DuoSet ELISA kits (R&D Systems). Data were acquired with a Molecular Device Spectra Max Plus 384 plate reader. Results were plotted using the Matplotlib library v3.10 (https://matplotlib.org). Statistics were calculated with the statistics functions in the SciPy library v1.15 (https://scipy.org).

### Co-immunoprecipitation

HEK293T cells were co-transfected with plasmids encoding full-length PA-tagged IL-17RA, Flag-tagged IL-17RB, and PA-tagged ACT1 using Lipofectamine 3000 (Invitrogen). At 24 hours post-transfection, cells were lysed in buffer (50 mM Tris-HCl, pH 7.5, 300 mM NaCl, 5% glycerol, 1% NP-40, 1% DDM, 0.2% CHS) supplemented with protease inhibitors (Roche) for 20 min on ice. Lysates were centrifuged at 14,000 g for 10 min at 4°C to remove debris. Supernatants were incubated with anti-Flag beads (Sigma-Aldrich) for 2.5 hours at 4°C with gentle rotation. Beads were washed three times with the lysis buffer and eluted by boiling in SDS loading buffer. Eluates were analyzed by SDS–PAGE followed by western blot. Anti-PA antibody (Fujifilm wako), Anti-Flag antibody (Sigma), Anti-mouse IgG (H+L) (DyLight 680 Conjugate)(Cat#5470S, cell signaling), and IRDye 800CW Goat anti-Rat IgG secondary antibody (LI-COR) were used. Inputs (4% of total) were loaded in parallel to for analyzing protein expression levels. The primary and secondary antibodies were used at 1:1000 and 1:10000 dilution, respectively.

### Negative-stain Electron Microscopy

Samples of different combinations of IL-17R fusions and ACT1 at 0.05 mg/ml was applied to glow-discharged copper grids (Electron Microscopy Sciences, # FCF400CU50) and incubated for 1 min. The grids were stained with 2% uranyl acetate for 1 min, blotted to near dryness with filter paper and air-dried. The grids were then stained with 2% uranyl acetate for 1 min and air-dried. Images were collected using a JEOL 1400+ electron microscope at 20000X magnification.

### Cryo-EM data collection and image processing

The IL-17RA/RB/ACT1 complex at 1 mg/ml was applied to glow-discharged Quantifoil R1.2/1.3 300-mesh gold holey carbon grids (Quantifoil, Micro Tools), blotted at 4 °C, 100% humidity and plunge-frozen in liquid ethane with a Mark IV vitrobot (Thermo Fisher). A total of 4536 Movies was collected in the super-resolution, correlated double sample mode on a Tritan Krios microscope (Thermo Fisher) with a K3 summit direct electron detector (Gatan). The slit of the GIF-Quantum energy filter was set to 20 eV. The nominal magnification of the microscope was set to 81,000, which corresponds to the pixel size of 1.07 Å. Movies were motion-corrected and dose-weighted by using MotionCor2 (v.1.2)^37^. Contrast transfer function (CTF) was estimated with GCTF1 1.06^38^.

Micrographs showed that the IL-17RA/RB/ACT1 complex formed filaments (Fig. 1b). Most of the filaments are around 300-400 Å long, but longer and shorter ones were also present. Filaments were picked from 3 micrographs using the napari_boxmanager in the crYOLO package (version 1.91)^39^. These filaments were used to train a model in the filament mode in crYOLO, which was then used for automatic filament picking from all micrographs. Filaments were segmented into overlapping boxes with an offset of 30 pixel in crYOLO and extracted into particles of box size of 200 pixel in RELION 5^40^. Although the filaments lacked true helical symmetry due to the presence of three different proteins, image processing in the helical reconstruction mode in RELION resulted in a reconstruction to 3.3 Å resolution (Supplementary Fig. 1). This was likely because multiple rounds of 3D classifications with imposed helical symmetry led to selection of segments that only contained ACT1 SEFIR domains but not those of IL-17RA or IL-17RB, which largely conform to one helical symmetry. Fourier-Bessel indexing of power spectra of the helical segments using the PyHI program suggested that the ACT1 filament has the helical symmetry of a one-start helix with 17.1 Å rise and 163.6° twist per subunit (Supplementary Fig. 1)^41^. This symmetry is equivalent to a two-start helix with 32.2 Å rise and 32.7° twist, but having handedness opposite to that of the one-start helix (Supplementary Fig. 1). The actual handedness of the helix was determined when the reconstruction reached high resolution that clearly resolved ⍺-helices in the proteins. These analyses showed that the ACT1 filaments could be described as a left-handed double-stranded helix. The refined rise and twist of the helical reconstruction are 16.8 Å and 162.9°, respectively, corresponding to the rise and twist of 33.6 Å and -34.2° respectively for the two-start helix.

To resolve IL-17RA, IL-17RB and ACT1 in the filaments, the data were reprocessed in the single-particle mode in RELION5 (Supplementary Fig. 2). Two-D class averages from the helical reconstruction above were used as templates to pick particles in the single particle mode. The minimal inter-particle distance was set to a small value (30 Å) to allow overlapping segments of filaments to be picked, analogous to picking of overlapping helical segments described above. A small box size (160 pixels) was used for initial particle extraction to reduce image processing time. The final box size was set to 400 pixels to maximize the chance of including full-length filaments that contained both IL-17RA and RB as well as multiple copies of ACT1. Well-ordered filament segments were selected through several rounds of 2D classifications, which are followed by multiple rounds of 3D classification steps. Reconstructions from the 3D classification and refinement showed that the SEFIR domain of IL-17RA has an extra ⍺-helix (⍺F) that is absent in both IL-7RB and ACT1. This unique feature allowed the selection from the results of 3D classification of particles that contained IL-17A, while excluding those that only contained ACT1. Particles were subjected to 3D refinement, CTF refinement and Bayesian polishing, leading to a reconstruction to 3.2 Å resolution (Supplementary Fig. 2). In this reconstruction, two IL-17RB and IL-17RA molecules constitute the first and second layers of the double-stranded filament, which are followed by multiple layers of ACT1 molecules. The two C-terminal tails form the two IL-17RA molecules extend along the filament axis to act as a tendril-like structure to help stabilize the ACT1 layers.

More careful inspection of the reconstruction revealed that the density for helix ⍺F in IL-17RA appeared much weaker than nearby ⍺-helices, suggesting misalignment that placed IL-17RB or ACT1 in some particles to the position of IL-17RA. A round of 3D classification without alignment confirmed the presence of the misalignment, which was evident from the different placements of the two IL-17RA molecules in the 3D class averages (Supplementary Fig. 2). The Euler angles and the origins of the particles in the misaligned classes were changed based on the rotations and shifts relative to the dominant class as shown in Supplementary Fig. 2. Particles with corrected rotation angles and origins were subjected to another round of 3D refinement with local search to prevent reoccurrence of misalignment. This was followed by CTF refinement and Bayesian polishing in RELION5, leading to a reconstruction of the overall resolution of 3.1 Å, with helix ⍺F in IL-17RA much better resolved. The density of the middle portion of the reconstruction was of high quality, but the two ends were blurred presumably due to bending motions along the axis of the filament. Improved density for the ends was obtained by two focused refinements with masks covering each end, respectively, which both reached 3.0 Å resolution. Two composite maps were generated from the two half maps from the three reconstructions in Phenix 1.9^42^. The two composite maps were combined into one map, with the FSC curve between the two maps calculated using the postprocessing routine in RELION. This combined composite map was used for atomic model building. The map contained two IL-17RB and two IL-17RA molecules, and seven and eight ACT1 subunits in the two strands respectively.

Atomic model building was initiated by docking the AlphaFold models of the SEFIR domains of *human* IL-17RA, RB and ACT1 into the composite map from the single-particle reconstruction in UCSF ChimeraX 1.7^24,43^. Atomic model was not built for the map from the helical reconstruction, because it does not contain more information than the single-particle reconstruction. In addition, the different β-sheet modules around different ACT1 subunits along the helix were averaged out by the imposed helical symmetry, leading to loss of information. The composite map from the single-particle reconstruction allowed unambiguous assignment of the three proteins into the density. The models were manually adjusted based on the densities, with the AlphaFold predicted model of the IL-17RA/RB/ACT1 complex as a reference. The model was first refined using the ISOLDE plugin (version 1.7) in ChimeraX^44^. Further real-space refinement of the coordinates with secondary structure restraints and atomic B-factors were carried out in Phenix 1.19. MolProbity was used for comprehensive validation of the models ^45^. Map resolution was estimated by using the gold-standard Fourier Shell Correlation criterion (cutoff at 0.143). The correlation between the map and model was calculated based on FSC correlation with the cutoff of 0.5. Figures were rendered in ChimeraX. The statistics of data collection and model refinement were summarized in Supplementary Table 1.

### AlphaFold Predictions

All the predictions were carried out by using the AlphaFold 3 server (https://alphafoldserver.com)^25^. For predicting individual SEFIR domains, the input sequences were IL-17RA (residues 379-588; Uniprot code: Q96F46-1), IL-17RB (residues 331-481; Uniprot code: Q9NRM6-1), IL17RC (residues 584-769; Uniprot code: Q8NAC3-1), IL-17RD (residues 356-566; Uniprot code: Q8NFM7) and IL-17RE (residues 488-667; Uniprot code: Q8NFR9-1). For the IL-17RA/co-receptor/ACT1 complexes, the inputs included two copies of IL-17RA (residues 361-866), 12 copies of ACT1 (residues 398-574; Uniprot code: O43734-1), and two copies of one of the co-receptors: IL-17RC (residues 581-776), IL-17RD (residues 351-510) and IL-17RE (residues 461-640). For predicting the IL-17RD1/ACT1 complex, two copies of IL-17RD (residues 341-739) and 8 copies of ACT1 (residues 398-574) were used. pIDDT (Predicted Intrinsic Distance Difference Test) scores and PAE (Predicted alignment errors) scores of the predictions are shown in Supplementary Fig 9.

### Data availability

The composite cryo-EM maps and the associated atomic model of the IL-17RA/RB/ACT1 complex structure have been deposited in the EMD (accession ID: EMD-70818) and RCSB databases (PDB ID: 9OT1), respectively. The consensus map, two focused refined maps used for generating the composite map and the map from the helical reconstruction of the ACT1 portion have been deposited as related entries in the EMD database with the accession IDs of EMD-70814, EMD-70815, EMD-70816, EMD-70817, respectively. The following previously reported structures were used for comparison: MyD88 (PDB ID: 7BEQ), the SEFIR domains of IL-17RA (PDB ID: 4NUX) and IL-17RB (PDB ID: 3VBC). Unless otherwise stated, all data supporting the results of this study can be found in the article, supplementary, and source data files. Source data are provided with this paper.

## Acknowledgements

Cryo-EM data were collected at the University of Texas Southwestern Medical Center (UTSW) Cryo-Electron Microscopy Facility, funded in part by the Cancer Prevention and Research Institute of Texas (CPRIT) Core Facility Support Award RP220582. We thank D. Stoddard and J. Martinez Diaz for facility access. We thank the Structural Biology Laboratory at UTSW for equipment use. This work is supported in part by grants from the National Institutes of Health (R35GM130289 to X.Z. and R35GM156386 to X.-c.B.), the Welch foundation (I-1702 to X.Z. and I-1944 to X.-c.B.). X.-c.B. and X.Z. are Virginia Murchison Linthicum Scholars in Medical Research at UTSW.

## Author contributions

H.Z., X.-c.B. and X.Z. conceived the project. H.Z. performed the biochemical and cell biological experiments. All authors contributed to cryo-EM data collection, structural determination and manuscript writing.

## Supplementary Figures

**Supplementary Figure 1.**
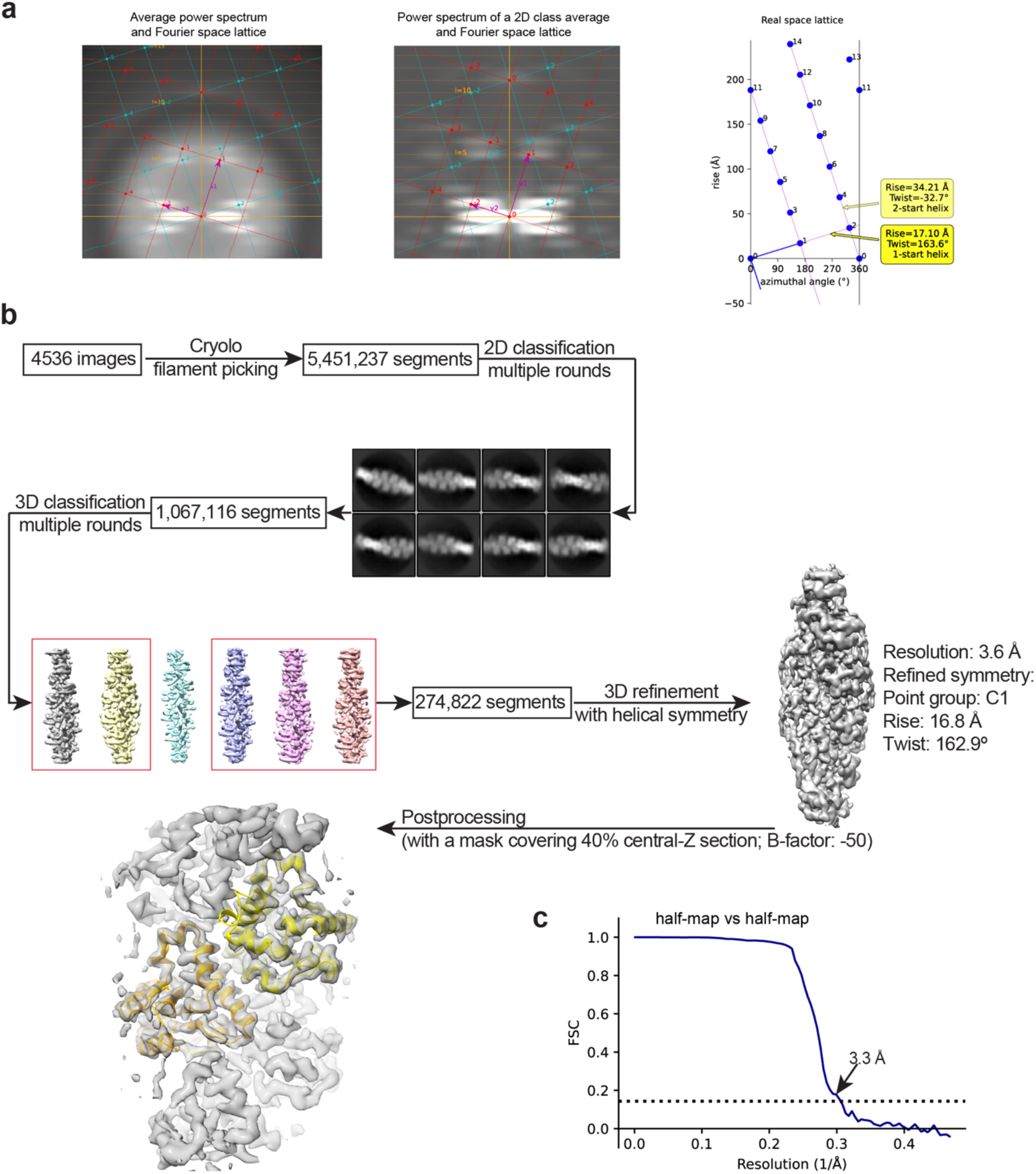
Helical reconstruction of the ACT1 portion of the IL-17RA/RB/ACT1 helical assembly. **a**, Deducing the helical rise and twist through Fourier-Bessel indexing of power spectra of the filaments. Results from indexing either an average power spectrum or the power spectrum of a 2D class average were essentially identical. **b**, Image processing process. **c**, Gold-standard FSC curve of the reconstruction.

**Supplementary Figure 2.**
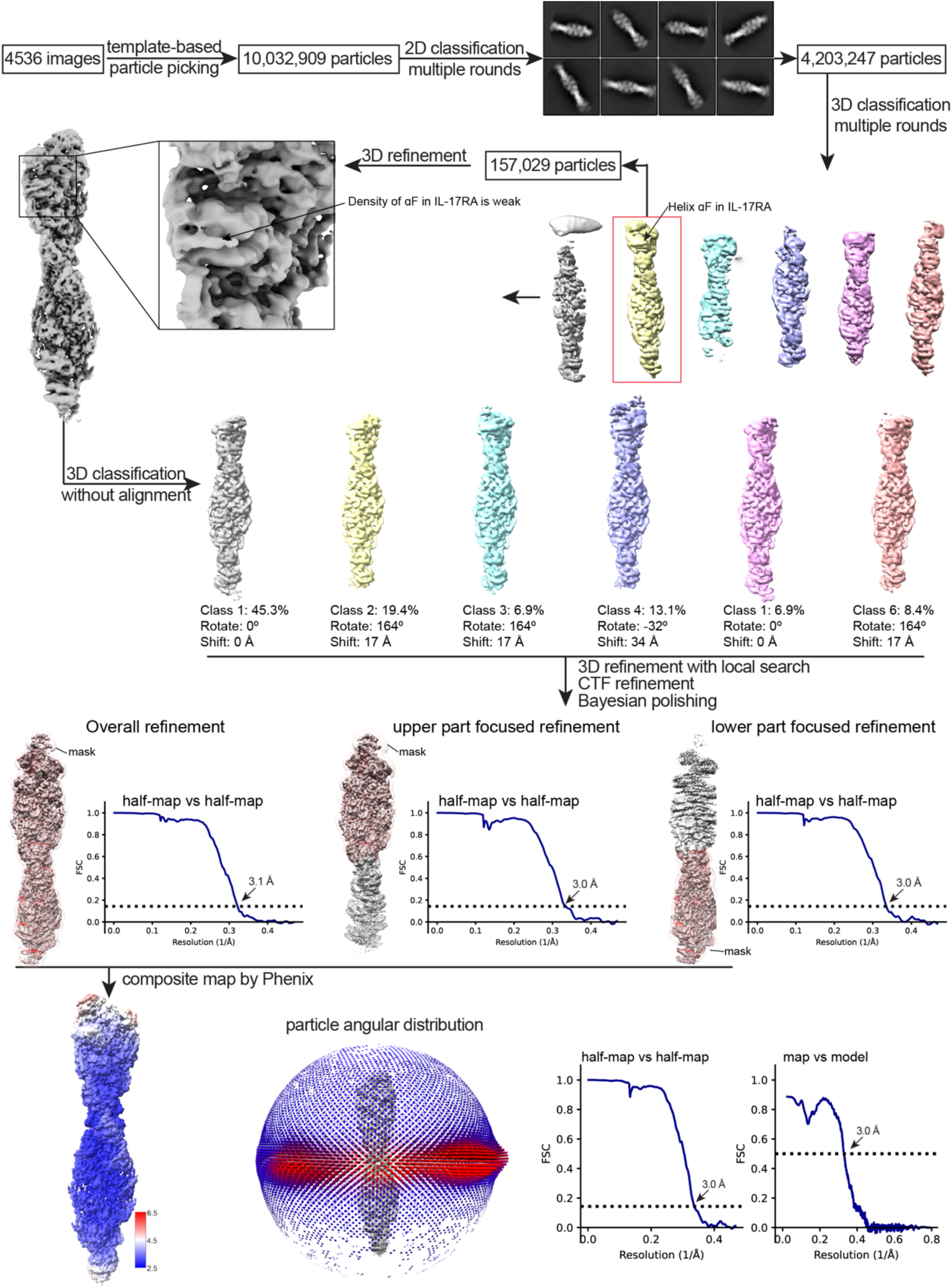
Single-particle reconstruction of the IL-17RA/RB/ACT1 complex.

**Supplementary Figure 3.**
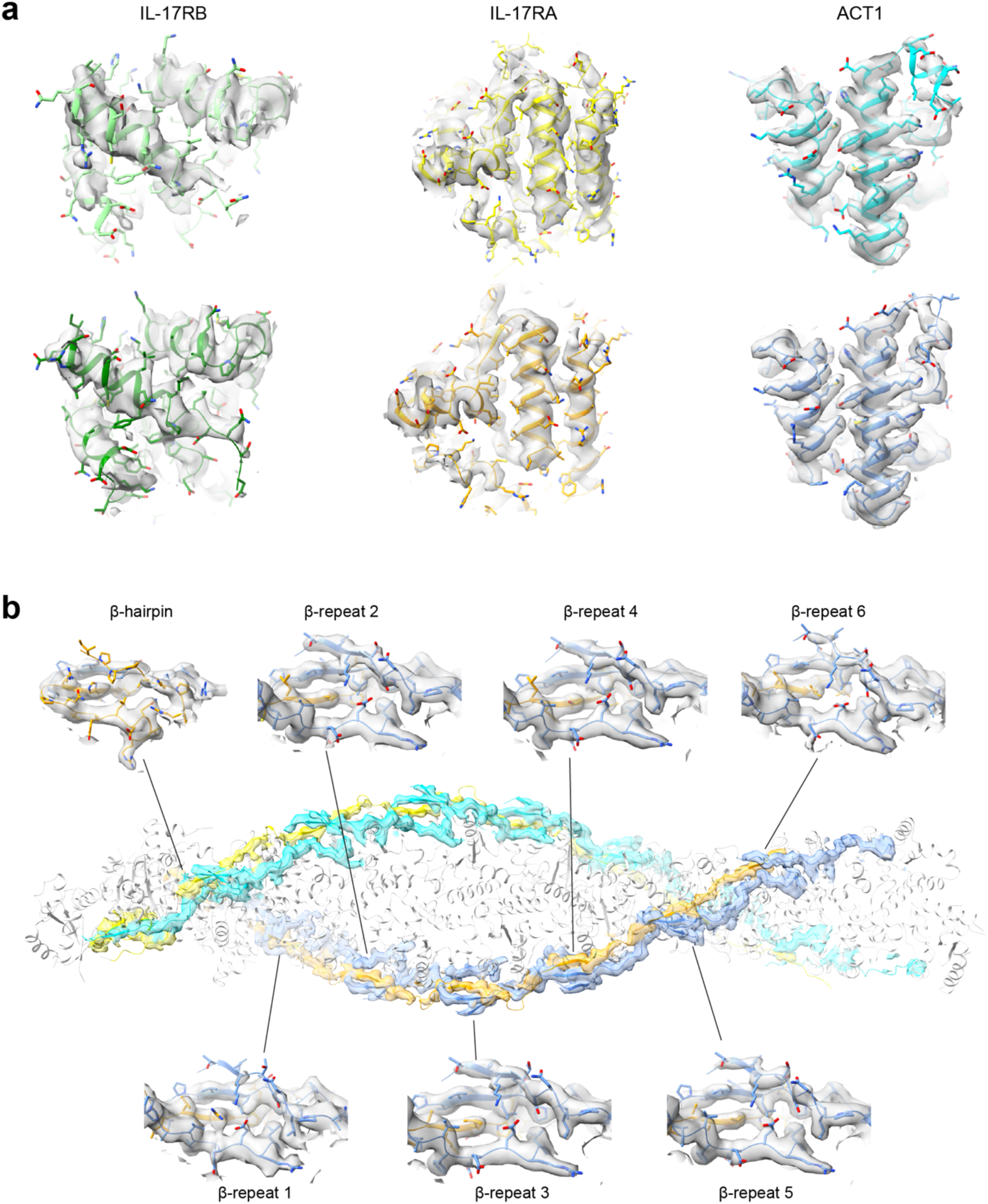
Samples of the cryo-EM density map. **a**, Density maps of the SEFIR domains of IL-17RB, RA and ACT1. **b**, Density maps of the β-hairpin and β-repeats.

**Supplementary Figure 4.**
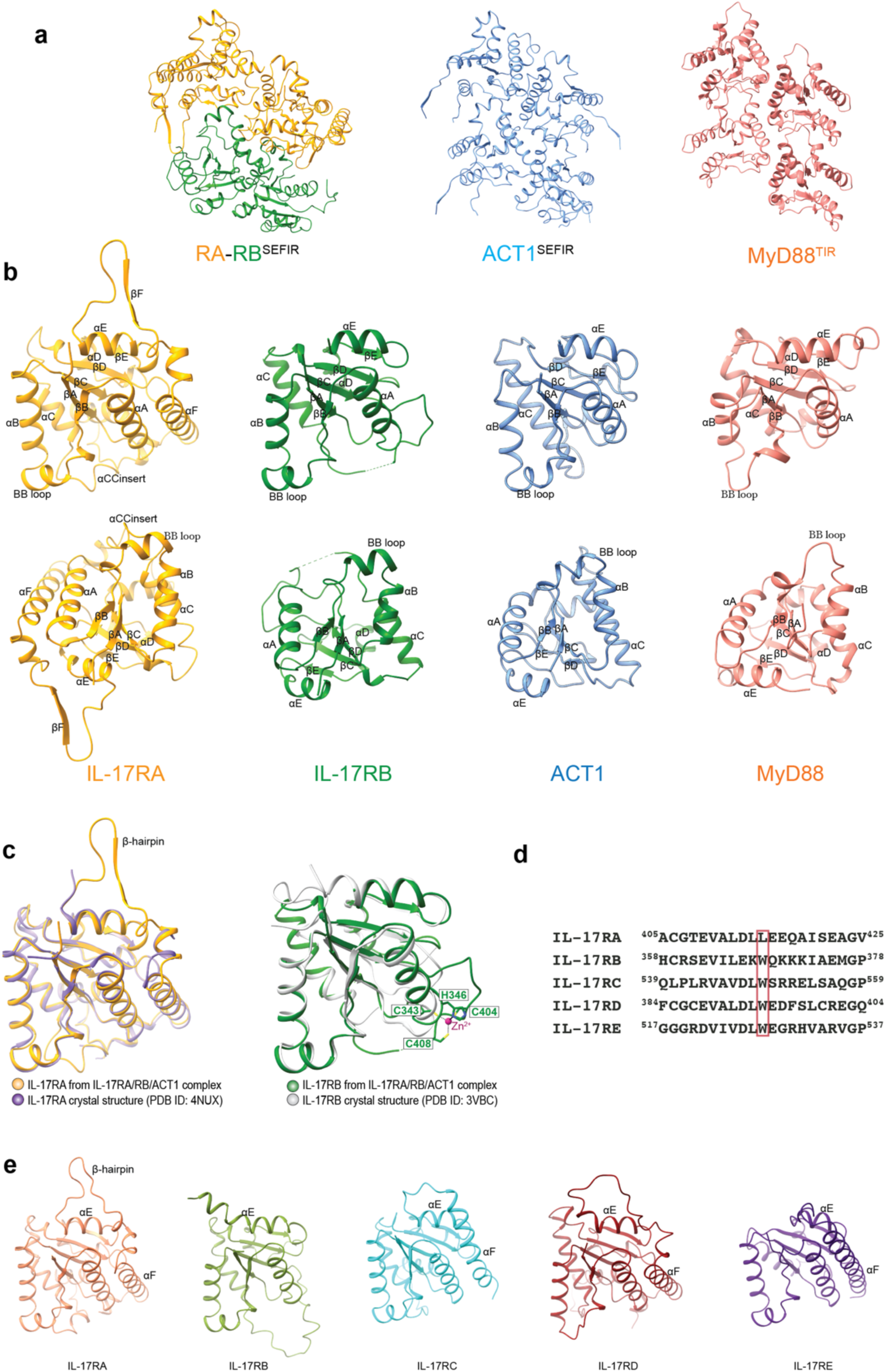
Comparison of SEFIR domains. **a**, Comparison of the assembly modes in the IL-17RA/RB/ACT1 complex and that of MyD88 TIR domain (PDB ID: 7BEQ). Four subunits are shown in each assembly. **b**, Comparison of the SEFIR domains of IL-17RA, RB, ACT1 in the cryo-EM structure and the TIR domain of MyD88 (PDB ID: 7BEQ). The structures are shown in two different orientations. The secondary structural elements are labelled. **c**, Comparisons of the SEFIR domains of IL-17RA and IL-17RB in the cryo-EM structure with previously published crystal structures. **d**, Sequence alignment showing that W368 in IL-17RB is conserved in other co-receptors but not in IL-17RA. **e**, Comparison of AlphaFold predicted models of the SEFIR domains of the IL-17Rs.

**Supplementary Figure 5.**
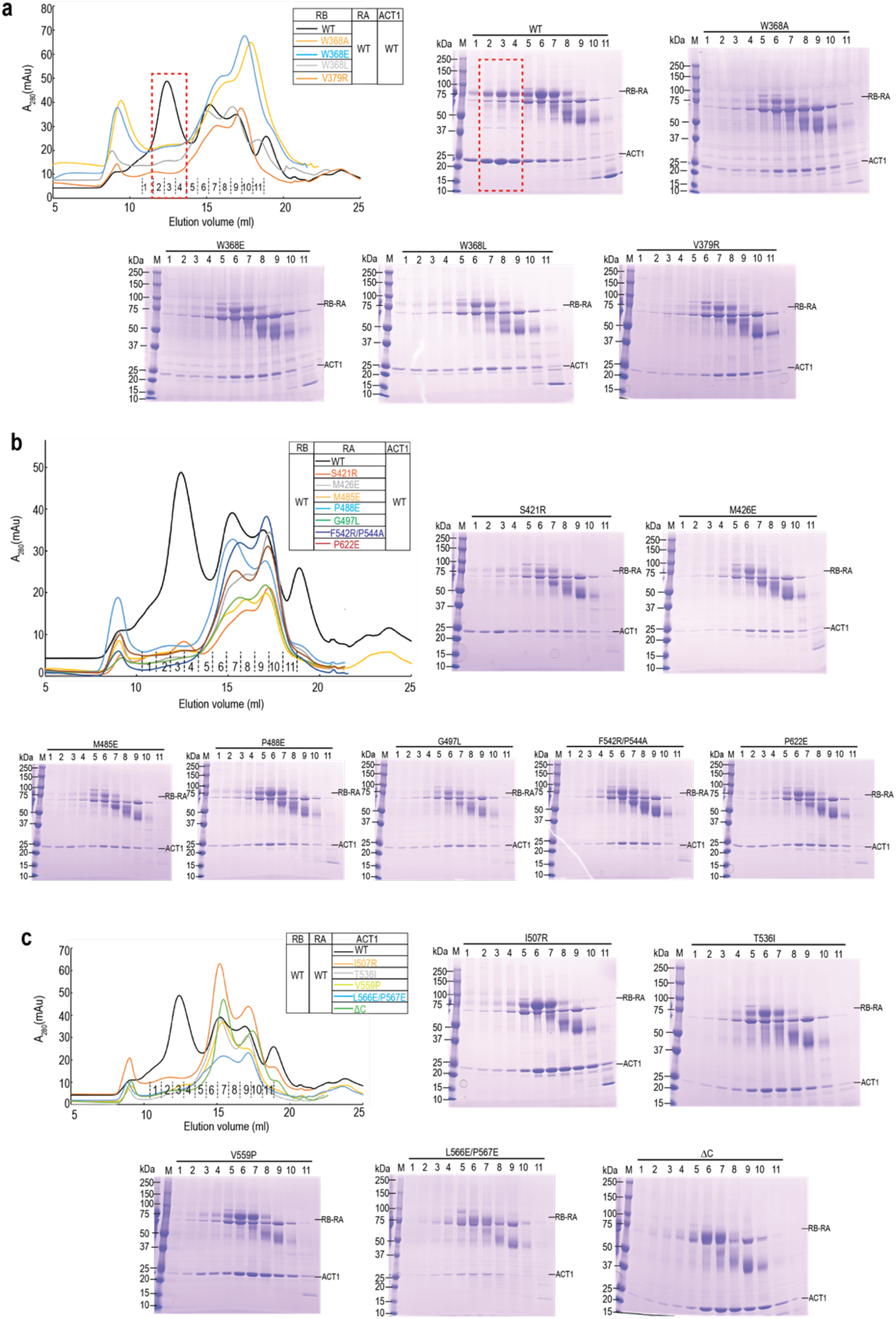
Gel filtration and SDS-PAGE analyses of the effects of the interface mutations on the formation of the IL-17RA/RB/ACT1 complex. **a**, Mutations of IL-17RB abolish the formation of the complex. The complex of wild type IL-17RA, RB and ACT1 is shown for comparison (fractions 2-4, red dashed boxes). **b**, Mutations of IL-17RA abolish the formation of the complex. **c**, Mutations of ACT1 abolish the formation of the complex.

**Supplementary Figure 6.**
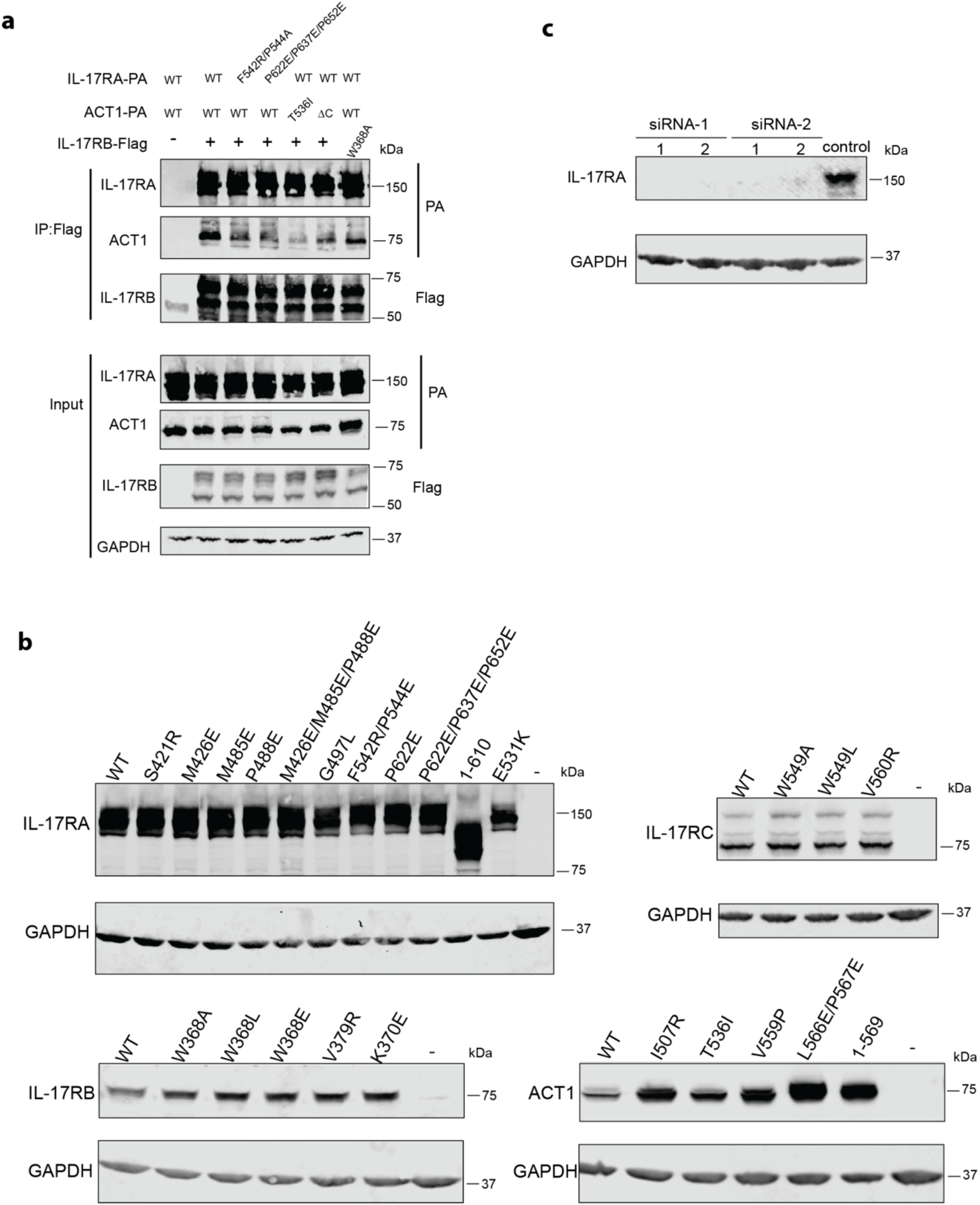
Western blot analyses of the expression and interaction of IL-17 receptors and ACT1. **a**, Interaction of ACT1 with IL-17RA, RB in HEK293T cells analyzed by co-immunoprecipitation. **b**, Western blot analysis of expression of the wild type and mutants of IL-17RA, RB, RC and ACT1 in HeLa cells. **c**, Western blot showing the knock-down effects of siRNA-1 and siRNA-2 targeting IL-17RA. The results shown are representative from at least three biological repeats.

**Supplementary Figure 7.**
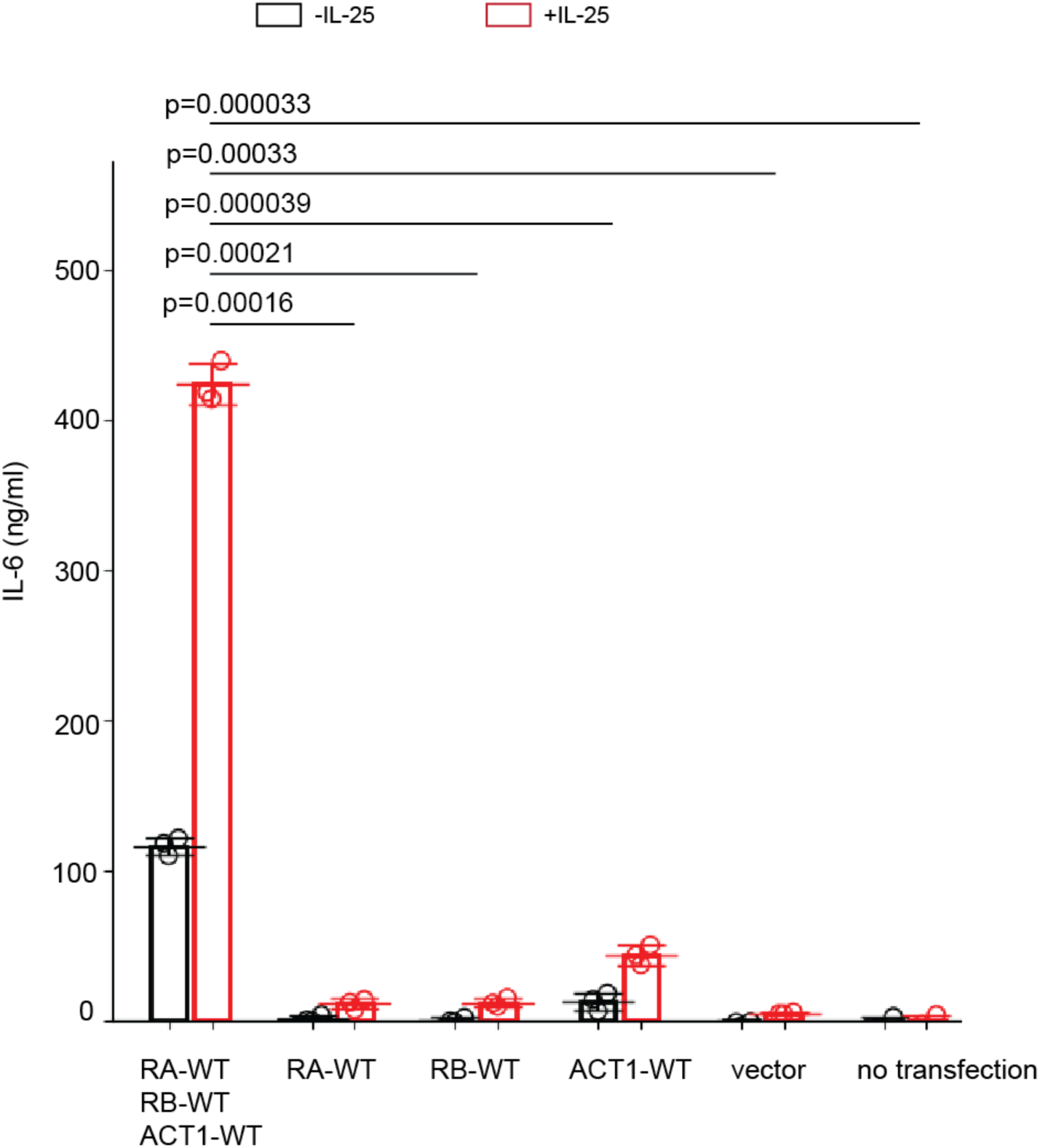
IL-25 induced robust production of IL-6 in Hela cells co-transfected with IL-17RA, IL-17RB, and ACT1. IL-16 secretion levels from un-transfected cells or cells transfected with an empty vector or only one of the three proteins were much lower. Each circle represents the result from one of three independent experiments. The bars represent means and standard deviations. P-values were calculated with two-tailed Student’s T-test.

**Supplementary Figure 8.**
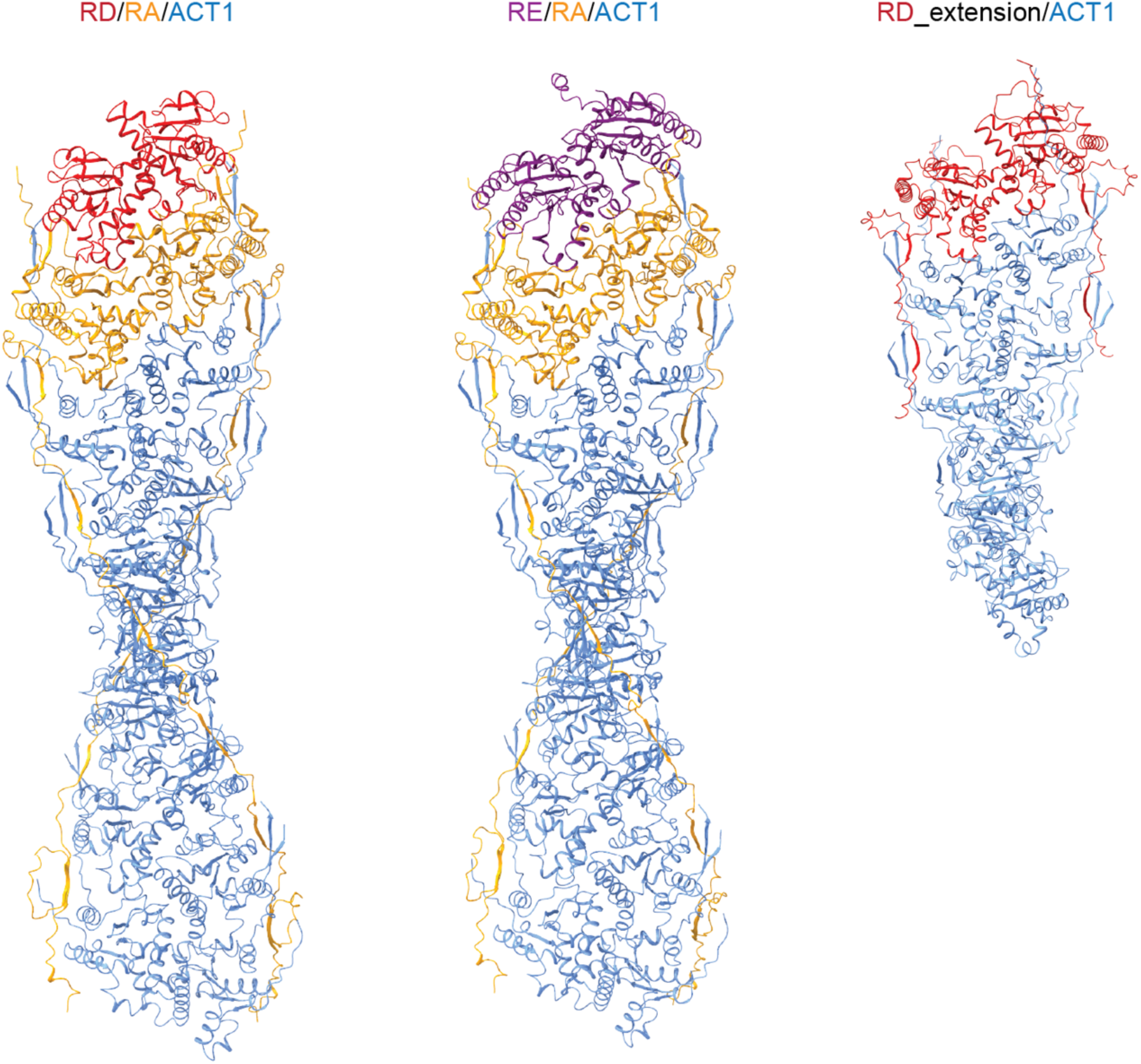
AlphaFold models of the helical assemblies of IL-17RA/RD/ACT1, IL-17RA/RE/ACT1 and IL-17RD/ACT1.

**Supplementary Figure 9.**
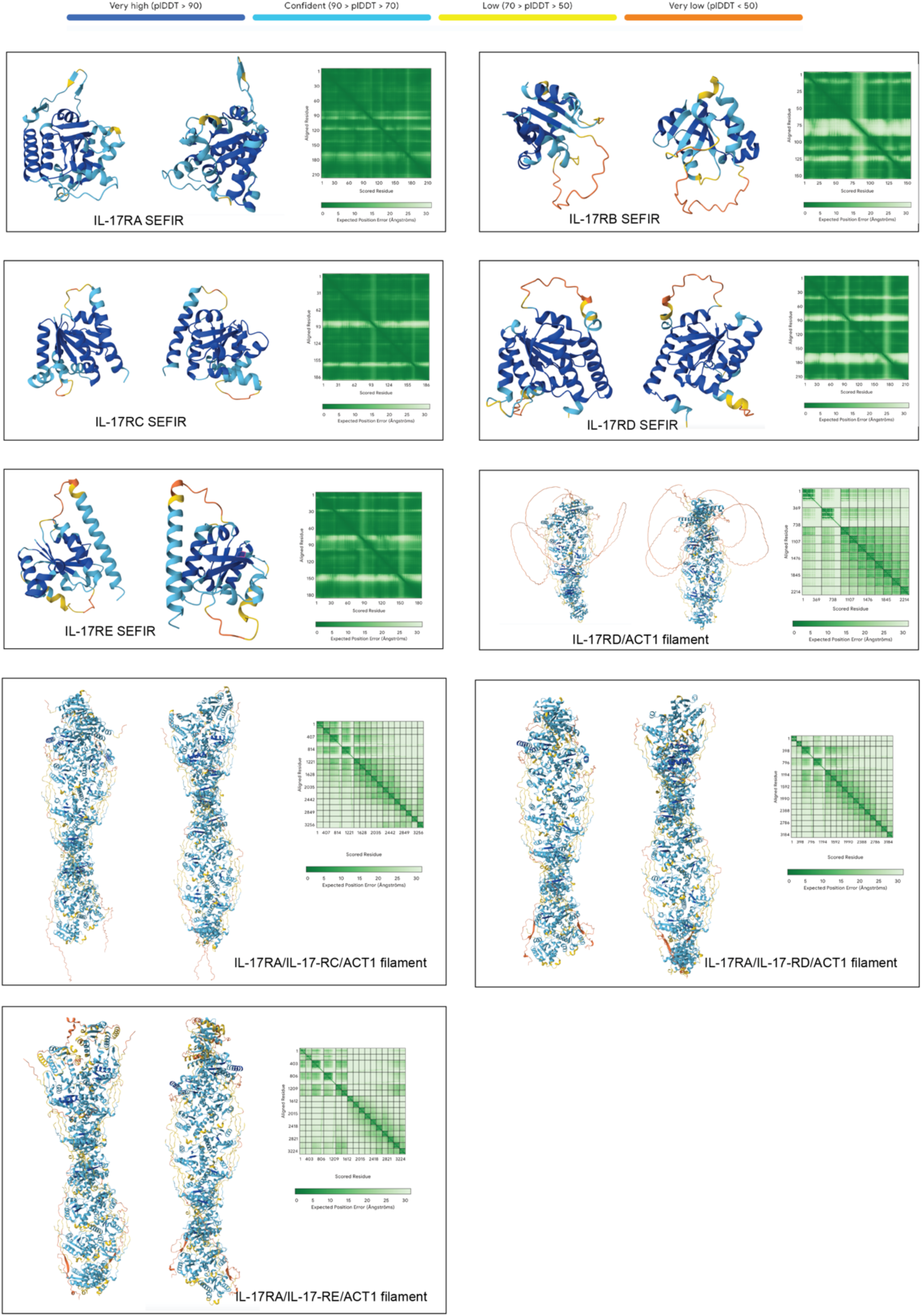
pIDDT (Predicted Intrinsic Distance Difference Test) scores and PAE (Predicted alignment errors) of the AlphaFold models shown in the paper.

## Supplementary Table

**Supplementary Table 1.** Cryo-EM data collection and structure refinement statistics.

|  |  |
| --- | --- |
| Magnification | 81,000 |
| Voltage (kV) | 300 |
| Electron exposure (e<sup>-</sup>/Å<sup>2</sup>) | 50 |
| Defocus range (μm) | 1.0-2.5 |
| Pixel size (Å) | 1.07 |
| Symmetry imposed | C1 |
| Initial particle images (no.) | 10,032,909 |
| Final particle images (no.) | 157,029 |
| Map resolution (Å) | 3.1 |
| FSC threshold | 0.143 |
| Initial model used<br>(AlphaFold model #) | IL-17RA: AF-Q96F46-F1<br>IL-17RB: AF-Q9NRM6-F1<br>ACT1: AF-O43734-F1 |
| Model resolution (Å) | 3.0 |
| FSC threshold | 0.5 |
| Map sharpening B factor (Å<sup>2</sup>) | 0 |
| Model Composition |  |
| Non-hydrogen atoms | 28,710 |
| Protein residues | 3510 |
| Ligands | 2 Zn<sup>2+</sup> |
| Protein B factors (Å<sup>2</sup>) | 110.2 |
| Ligand B factors (Å<sup>2</sup>) | 273.0 |
| R.m.s. deviations |  |
| Bond length (Å) | 0.004 |
| Bond angle (°) | 0.53 |
| Validation |  |
| Molprobity score | 1.2 |
| Clashscore | 3.1 |
| Poor rotamers (%) | 0.2 |
| Ramachandran plot |  |
| Favored (%) | 97.5 |
| Allowed (%) | 2.5 |
| Outliers (%) | 0 |

